# From prior information to saccade selection: evolution of frontal eye field activity during natural scene search

**DOI:** 10.1101/251835

**Authors:** Joshua I. Glaser, Daniel K. Wood, Patrick N. Lawlor, Mark A. Segraves, Konrad P. Kording

**Affiliations:** Interdepartmental Neuroscience Program, Northwestern University, Chicago, IL, USA; Department of Physical Medicine and Rehabilitation, Northwestern University and Shirley Ryan Ability Lab, Chicago, IL, USA; Department of Bioengineering, University of Pennsylvania, Philadelphia, IL, USA; Department of Neurobiology, Northwestern University, Evanston, IL, USA; Division of Neurology, Children’s Hospital of Philadelphia, Philadelphia, PA, USA; Department of Neuroscience, University of Pennsylvania, Philadelphia, IL, USA

## Abstract

Prior knowledge about our environment influences our actions. How does this knowledge evolve into a final action plan and how does the brain represent this? Here, we investigated this question in the monkey oculomotor system during self-guided search of natural scenes. In the frontal eye field (FEF), we found a subset of neurons, “early neurons,” that contain information about the upcoming saccade long before it is executed, often before the previous saccade had even ended. Crucially, much of this early information did not relate to the actual saccade that would eventually be selected. Rather, it related to prior information about the probabilities of possible upcoming saccades based on the pre-saccade fixation location. Nearer to the time of saccade onset, a greater proportion of these neurons’ activities related to the saccade selection, although prior information continued to influence activity throughout. A separate subset of FEF neurons, “late neurons”, only represented the final action plan near saccade onset and not prior information. Our results demonstrate how, across the population of FEF neurons, prior information evolves into definitive saccade plans.

## Introduction

Deciding where to look next in the real world is a complex process, as we must rapidly decide between countless options. Prior knowledge about the environment and past behavior can facilitate decisions by focusing finite computational resources on options that have a higher probability of success. For example, if you are currently looking on the left side of the desk for a pencil, it will be most useful to look rightwards next. Using prior information to make preliminary plans about upcoming saccades could be an efficient use of neural resources in the oculomotor system.

To be more precise, in visual search, we define prior information about the upcoming saccade as anything that can influence the saccade decision before the subject has access to new visual information. In a Bayesian framework, we can think of the new visual information as the “likelihood,” and this likelihood is combined with the prior information to make a saccade decision In classic neuroscience visual search tasks (e.g. [1,2]), where stimuli are flashed onto the screen, prior information can be any knowledge or biases that can affect the upcoming saccade prior to the stimuli being displayed. In more natural visual foraging (e.g. [3,4]), where a subject is making continuous saccades around a scene, prior information is anything that could affect saccade planning prior to processing the new visual information at each new fixation location. This could include general information about the task or environment, information gathered during previous saccades, search strategies and biases, and more. There are likely many ways in which prior information could affect saccade planning.

Several previous studies have shown that oculomotor structures in the brain utilize prior information for planning saccades [1,2,5–9]. In macaque superior colliculus (SC), neurons show increased pre-target activity [5,6] when there is an increased probability that a target will be placed in the neurons’ receptive fields. For SC neurons [9] as well as for frontal eye field (FEF) corticotectal neurons [7], prior information about the task (whether it is a pro-saccade or anti-saccade task) affects pre-target activity. Additionally, the identity of targets on previous trials can affect the response of FEF neurons to targets on the current trial [1,2]. Thus, there is evidence that prior information affects neural activity at cortical and midbrain levels of the oculomotor system.

However, unlike unconstrained, natural eye movement behavior, most tasks used in previous studies imposed substantial limitations on the available prior information. Rather than eliciting self-guided search behavior, these tasks elicited single saccades instructed by a target. This approach eliminated the ongoing planning of sequences of saccades, which is a function of FEF neurons in natural search conditions [4,10]. For example, monkeys might carry over upcoming saccade plans from previous saccades. Conventional tasks also remove the possibility of ruling out saccade targets based on previous saccades [8]. Finally, these tasks often removed starting eye positions as a variable. The oculomotor system is modulated by eye position in a manner that favors movement towards the center of the oculomotor range [11]. Thus, by ignoring eye position, these previous studies also removed a significant source of prior information for constraining the range of potential eye movements. In naturalistic settings, much therefore remains unknown about how the oculomotor system represents prior information, and how this representation evolves into a definitive saccade plan.

Here, to explore how prior information affects saccade planning in more naturalistic conditions, we recorded from macaque FEF during a natural scene search task. A subset of neurons, “early neurons,” reflected the probabilities of upcoming eye movements based on the current eye position, regardless of the actual selected saccade direction. As time elapsed toward the upcoming saccade, the activity of these neurons began to relate more to the impending saccade direction, although the prior information continued to influence activity throughout. There was another subset of neurons, “late neurons” that only coded for the selected action plan shortly before the upcoming saccade, and did not represent prior information. Thus, across the population of FEF neurons, we observe how prior information evolves into definitive saccade plans.

## Results

### Behavior

To better understand the evolution of saccade plans during self-guided eye movements, we recorded single units from the frontal eye field (FEF) while head-fixed monkeys freely searched for a target embedded in natural scenes (Fig. 1A) [12,13]. Trials either ended when the monkeys made 20 saccades without finding the target, or when they made a saccade to the target and held gaze there to receive a reward. During such a self-guided search, monkeys could use prior information to start planning saccades before they have new detailed visual information at each upcoming fixation location.

**Figure 1:**
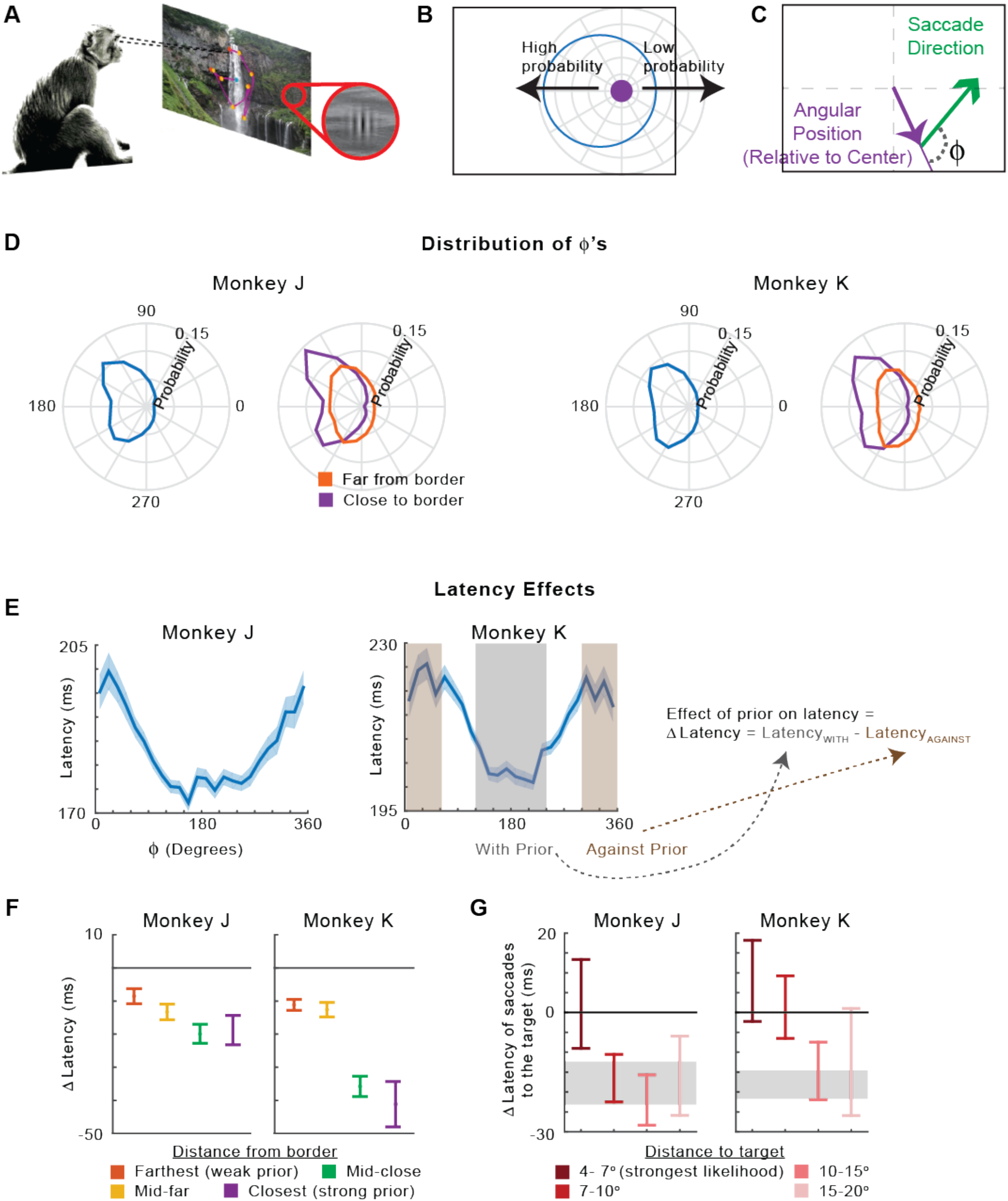
Experiment and behavior. **(A)** Monkeys freely searched for a Gabor target embedded in natural scenes. **(B)** The probability of the direction of the upcoming saccade is dependent on the eye position on the screen. This is an example where the eye position is to the right of the screen. **(C)** We quantify the relationship between the upcoming saccade direction and eye position using Φ, the angle between the angular position (the eye position vector relative to center), and the upcoming saccade vector. **(D)** The distribution of Φ’s for all saccades (blue), and split according to whether the starting eye position was close (purple) or far (orange) from the border. The close/far distinction was based on being less/more than the median distance from a border. **(E)** The mean latency of saccades as a function of Φ. Saccades back towards the center (“with prior”) have |Φ-180°|<60°. Saccades away from the center (“against prior”) have |Φ-180°|>120°. We use Δ *Latency* (the difference between latencies with and against the prior) as a behavioral metric of the effect of the prior. **(F)** Δ *Latency* as a function of the starting eye position’s distance from a border. The distance from the border was divided into quartiles. **(G)** Δ *Latency* for saccades that end near the target, as a function of the initial distance to the target. The shaded area represents the mean +/− SEM of the latency difference for saccades not to the target. In panels E-G, all error bars represent SEMs.

One easily quantifiable factor that could provide prior information is the eye position on the screen. For instance, when the monkey is fixating on the right side of the screen, there are more possible saccadic opportunities to the left, and thus the monkey might make preliminary plans to go left (Fig. 1B).

To explore this idea, we defined a quantity Φ, which was the angle between the angular eye position (the eye position vector relative to the center of the screen) and the upcoming saccade vector (Fig. 1C). When going back towards the center, Φ = 180°, and when going away from the center, Φ = 0°. We found that monkeys are more likely to look approximately opposite of their current angular position (away from the borders of the screen), and the effect is stronger when closer to the border of the screen; Fig. 1D). This is in line with the known finding of center bias in eye movement behavior [14–16]. Interestingly, the peak of Φ is not at exactly 180° (i.e., going back towards the exact center). In both monkeys, there is a higher probability of Φ = 135° or Φ = 225° than Φ = 180°. This is because these statistics do not simply reflect the on-screen saccades that are possible (which would be centered on 180°); they also reflect any other strategies and biases of the monkeys.

If prior information based on eye position matters for saccade planning, we would expect higher probability saccades to have shorter latencies. This was the case; latencies were shorter for saccades made approximately opposite of the angular eye position (at Φ close to 180°; both monkeys, *p* < 1e-10; Fig. 1E; see Fig. S1 for the distribution of all latencies). This finding is consistent with several previous studies showing that saccades back towards the center have shorter latencies [11,17,18].

We used this finding to create a behavioral metric for the effect of the prior on behavior. We defined Δ *Latency* as the latency difference between saccades going “with” the prior (Φ close to 180°) and saccades “against” the prior (Φ far from 180°; Fig. 1E). If the prior is having a stronger effect on behavior, then there should be a larger magnitude latency difference. Indeed, for eye positions closer to the border, when prior information was more informative, the magnitude of Δ *Latency* was larger (Monkey J, *p*=4.9e-4; Monkey K, *p*=7.9e-10; Fig. 1F). Overall, the monkeys’ behaviors suggest that the oculomotor system became more prepared to look in a given direction as the probability of a saccade in that direction increased.

We can also analyze the behavior from a Bayesian perspective, in which prior information is combined with a likelihood (here, the new visual information) to form a decision. When there is strong likelihood information, the prior information should have less influence on the final decision. In our task, when the next saccade is to the target, we can assume that the likelihood was generally very informative, as visual information about the target is driving the decision. Thus, when the next saccade is to the target, we would expect prior information related to eye position to be less influential.

To test how likelihood information modified the influence of prior information, we again used the Δ *Latency* metric. For saccades that went to the target, we plotted Δ *Latency* as a function of the distance to the target. We separated saccades based on the distance to the target, because it is likely that long saccades that landed near the target may have landed there by accident, rather than based on visual information (see Fig. S2 for probabilities of making a saccade to the target as a function of distance). We found that for shorter saccades to the target, there was not a significant latency difference between saccades towards and away from the center, suggesting the prior had a limited influence (Fig. 1G). For longer saccades to the target (>7° for Monkey J and >10° for K), there was a latency difference comparable to saccades not going to the target. This provided behavioral evidence that when there is a strong likelihood (a target nearby that is noticed), position-based prior information is less influential.

### Overview of Neural Data Analysis

Next, we analyzed the activity of single FEF neurons while the monkeys performed the natural scene search task. As we were aiming to understand the neural correlates of prior information, we initially focused on activity from around the time of fixation, before new visual information could be gathered. We thus identified “Early” neurons, which were already predictive of the upcoming saccade around the time of fixation. We then determined whether this prior-related activity was based on position in the manner expected by our behavioral results.

We were not only interested in how neural activity related to prior information, but also how it related to the final saccade selection. To disentangle between activity related to prior information and saccade selection, we used models that separated activity predicted by position (which was the basis of the prior information) and the actual upcoming saccade. Using these models, we determined how neural activity’s relation to prior information and saccade selection evolved as time elapsed from fixation to saccade onset. Additionally, as in our behavioral analysis, we took a Bayesian approach to determine the scenarios when early neural activity predicted the final saccade. Finally, we identified “Late” neurons, which were only predictive of the final selected saccade near the time of saccade onset, and not prior information. We have thus shown how natural search behavior can be viewed through the lens of Bayesian integration theory, and have mapped it onto subsets of neurons in the FEF.

### Early saccade-related neural activity

To investigate the neural basis of how prior information is used for saccade planning, we first looked at the time at which neurons’ activities began to be informative of the upcoming saccades. We used a generalized linear model (GLM)-based approach (see *Methods*) to determine how important the upcoming saccade vector was for predicting neural activity, beyond the effects of the previous saccade. We found that 38/180 (21%) of recorded neurons had activity that was significantly modulated by the upcoming saccade in the 50 ms around fixation (before new visual information could be processed), and we classified these as “Early neurons”.

We investigated these neurons with peri-event time histograms (PETHs) that compared neural activity prior to saccades toward the neurons’ preferred directions (PDs) versus away from the PDs (Figs. 2A and S3A). When looking at individual neurons’ responses (Fig. 2, rows 1-3) and the average across Early neurons (Fig. 2, bottom row, and Fig. S3), it was clear that the activity traces began to differentiate early, even before fixation onset. As Early neurons had predictive activity that preceded new visual information, this demonstrates that those neurons represented prior information about upcoming saccades.

**Figure 2:**
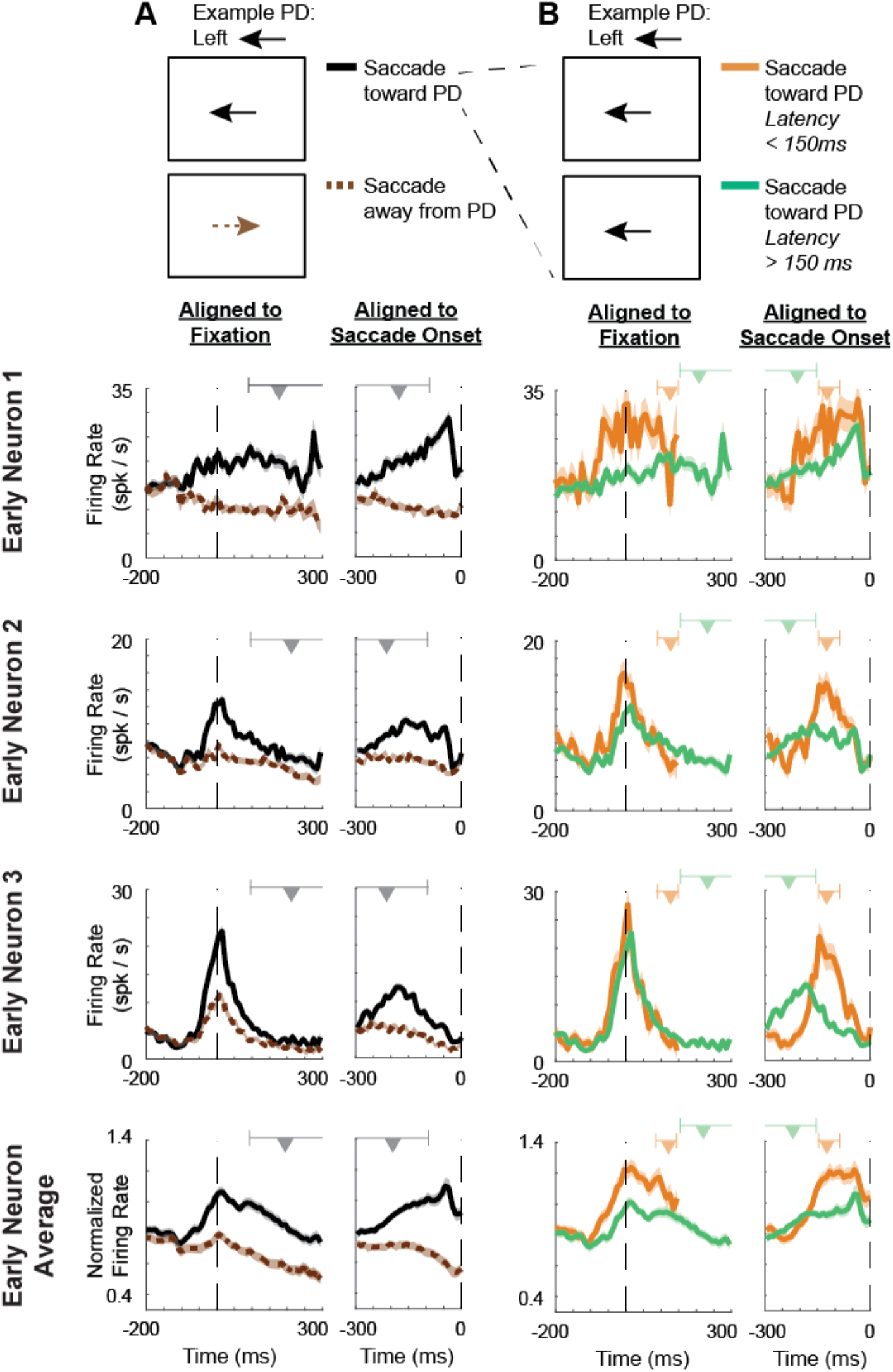
Early times of saccade selectivity, and neural differences related to saccade latencies. Peri-event time histograms (PETHs), aligned both to fixation (left part of each column) and the upcoming saccade onset (right part). **Rows 1-3 of PETHs**: Example Early neurons. **Bottom Row**: Normalized averages of all Early neurons. **(A)** PETHs of saccades toward the preferred direction (PD; black, solid) versus away from the PD (brown, dashed). Above the PETHs aligned to fixation, we show the range of 95% of saccade initiation times (the upper end of this range is larger than the x-axis limit). Above the PETHs aligned to saccade onset, we show the range of 95% of fixation onset times (the lower end of the range is below the x-axis limit). The triangles represent the median times. **(B)** PETHs of saccades toward the PD (like the black trace in panel A), divided further based on saccade latency. Saccades with latencies less than 150 ms are shown in orange while saccades with latencies greater than 150 ms are in green. Above the PETHs, ranges of fixation/saccade times are shown separately for the separate traces. In all figures, for the plots aligned to fixation, only data obtained before the onset of the saccade are included in the PETHs.

If the Early neurons’ activities are involved in saccade planning, then we would expect them to relate to the latency of the upcoming saccade, with higher activity predictive of shorter saccade latency [19–22]. We found that this is the case; for saccades into neurons’ PDs, shorter saccade latencies were associated with greater neural activity (Figs. 2B and S3B). This provides evidence that the prior information represented by Early neurons reflects aspects of the saccade-planning process.

### Prior information based on eye position

Our behavioral results suggested that eye position provides prior information that affects the planning process. Does the prior information encoded by Early neurons relate to the monkey’s eye position on the screen? We again used a GLM-based approach to determine whether eye position significantly modulated neurons’ activities. This model included eye position and the upcoming saccade (along with the previous saccade), so we could determine the influence of eye position beyond the saccade that actually occurs. We found that 28/38 (74%) of Early neurons were significantly modulated by position in the 50 ms around fixation. We will refer to these neurons as “Early/Pos” neurons. Thus, position may be used as a source of prior information within many Early neurons.

Since many Early neurons are modulated by eye position, it is possible that they use eye position as prior information to determine which saccades are possible or likely from that position. For instance, when the monkey is fixating on the left side of the screen, it may make preliminary plans to move rightward. Thus, if a neuron with a PD to the right was representing prior information about potential upcoming saccades, we would expect this neuron to have greater activity when the monkey’s eye position was on the left side of the screen. Generalizing this example, we would expect Early neurons to have greater activity when the monkey’s eye positions are such that the next saccade is likely to be into the neurons’ PDs. This occurs when the angular eye positions are approximately opposite of the neurons’ PDs. Importantly, we should be able to see evidence of this prior information based on position, regardless of the direction of the actual saccade. That is, we want to determine whether the prior information has its own independent influence upon activity.

Using PETHs, we investigated how the activity of Early/Pos neurons depended on the initial eye position. Indeed, we found that these neurons did have greater activity when the angular eye position at fixation was opposite of the neurons’ PDs (Figs. 3 and S4; red trace higher than blue trace), i.e., when the upcoming saccade is more likely to go into the neurons’ PDs. This differentiation of activity based on the initial eye position began prior to fixation, long before the upcoming saccade. This is around the same time these neurons became predictive of the upcoming saccade (Figs. 2 and S3). Importantly, even when controlling for the direction of the upcoming saccade, Early/Pos neurons still had greater activity when the angular eye position was opposite the PD (Figs. 3 and S4). In other words, neurons had higher activity when the eye was in a position that made a saccade into the PD likely, even if the saccade didn’t actually end up going into the PD (Figs. 3B and S4B). This finding, along with the early timing of these neurons’ responses, supports the idea that Early neurons use eye position as a source of prior information to make initial saccade plans.

**Figure 3:**
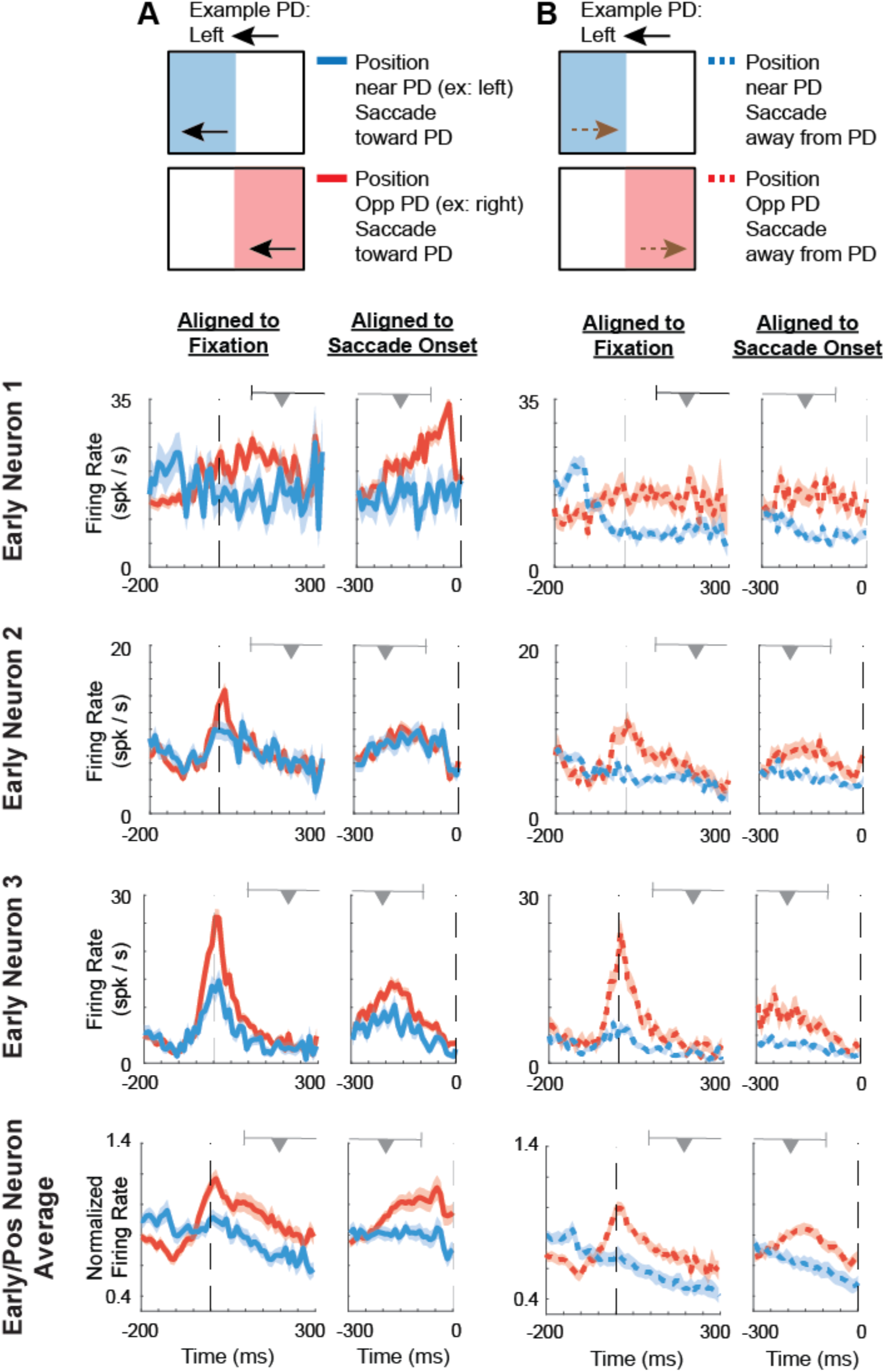
Increased activity in positions that are more likely to result in saccades toward the PD. Peri-event time histograms (PETHs), aligned both to fixation (left part of each column) and the upcoming saccade onset (right part of each column). Ranges of saccade/fixation onset times are shown above the PETHs as in Fig. 2. **Rows 1-3 of PETHs**: Example Early/Pos neurons. **Bottom Row**: Normalized averages of all Early/Pos neurons. **(A)** PETHs of saccades toward the PD, with a starting angular eye position near the PD (unlikely that upcoming saccade will be toward PD; blue) versus an angular position opposite the PD (likely that upcoming saccade will be toward PD; red). For example, if the PD is to the left, positions near the PD will be on the left side of the screen (see *Methods* for details). **(B)** PETHs of saccades away from the PD, with a starting angular position near the PD (blue, dashed) versus an angular position opposite the PD (red, dashed).

Beyond looking at when the angular eye positions are near vs. opposite neurons’ PDs, we want to understand how neural activity more specifically relates to positions across the screen. Does this activity relate to the probabilities of saccades that occur from a given position (Fig. 1D)? To investigate this, we tracked the average activity over time, as a function of the “relative angular position”. The relative angular position is the angular position relative to the PD of the neuron (Fig. 4A), and is an analog to the behavioral measure Φ (Fig. 1C, Fig. 4B). Importantly, we only included fixation periods preceding saccades that were made away from the neurons’ PDs in order to minimize contamination by neural activity that was related to the actual saccade itself.

**Figure 4:**
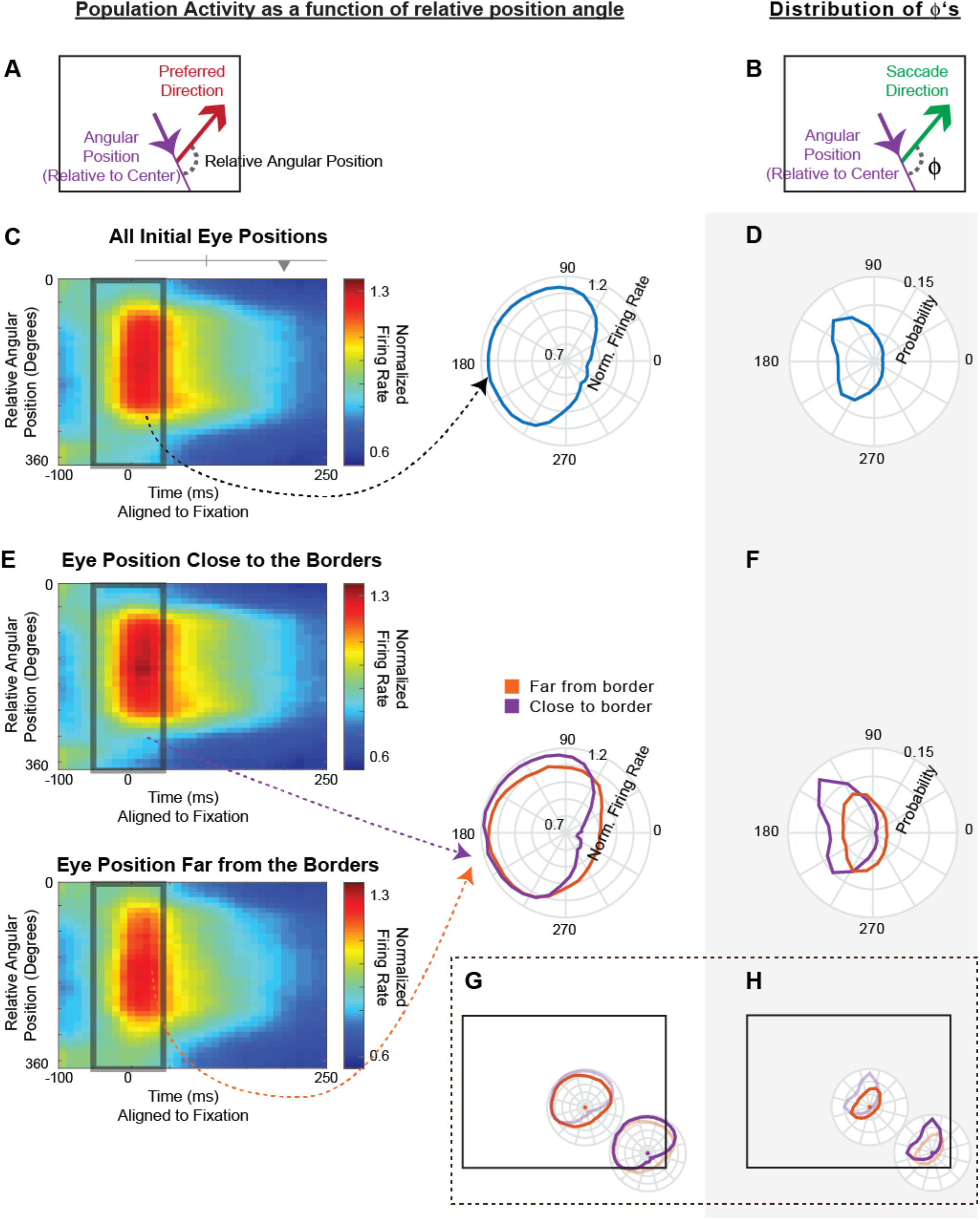
Early population activity reflects the probabilities of upcoming saccades. On the left, we plot the population activity of Early/Pos neurons as a function of the relative angular position. On the right, with a gray background, we show how this relates to the behavioral distribution of Φ’s. **(A)** The relative angular position is the difference between a neuron’s preferred direction and the angular eye position (the vector from the screen center). **(B)** Copied from Fig. 1c, Φ is the difference between the upcoming saccade and the angular eye position. **(C)** On the left, a heat map of normalized activity over time, as a function of relative angular position, averaged across neurons. Ranges of saccade onset times are shown above, as in Fig. 2. On the right, the normalized average activity in the 100 ms surrounding fixation, plotted as a function of the relative angular position. Only saccades away from the PD are included. **(D)** The distribution of Φ’s across all saccades combined across monkeys. **(E)** Same as panel C, but now separated for initial eye positions close to the borders (top left, right in purple) and far from the borders (bottom left, right in orange). Only saccades away from the PD are included. **(F)** The distribution of Φ’s combined across monkeys, separated by initial eye positions close to the borders (purple) and far from the borders (orange). **(G,H)** The distributions from panels E and F, respectively, are plotted on set positions, rather than as a function of relative angular position. The distribution corresponding to the set position (e.g. purple when close to the border) is darker.

In the resulting plot, we observed greater activity when the angular eye position was approximately opposite the neurons’ PDs (Figs. 4C and S5A), which agreed with our PETH results. We ran a control to ensure that this relation was not due to activity from the previous saccade (Fig. S5C). By looking more closely at the 100 ms around fixation, we found that the distribution of activity was generally similar to the behavioral distribution of Φ’s, although the activity distribution did not have the peaks at 135° and 225° seen in the behavior.

As the probability distributions of upcoming saccades depended on whether the eye position was close versus far from the border (Fig. 1D), we compared activity between these conditions in the same manner as above. We found that for angular eye positions opposite to the PD, there was greater activity when close to the border (Figs. 4E and S5B). This relates to the behavior, in that the probability of saccades opposite the current angular position is greater when closer to the border (Fig. 4F). For angular eye positions in the same direction as the PD, there was lower activity when close to the border (Figs. 4E and S5B). This relates to the behavior, in that the probability of saccades in the same direction as the current angular position is smaller when closer to the border (Fig. 4F). Moreover, by plotting the distributions of activity on given positions close or far from the borders (Fig. 4G,H), we can more intuitively see how the activity represents a prior distribution of potential movements from a given starting position. There is a wider distribution when farther from the border, and a narrower distribution when closer to the border. Thus, the early activity of the subset of Early/Pos neurons closely relates to the probabilities of saccades that will later occur.

### Evolution of neural activity over time

We have seen that, early on, the activity of Early/Pos neurons represents prior information about potential upcoming saccades. How does the neural activity evolve over time as the upcoming saccade approaches? We first looked at PETHs preceding saccades with latencies in small windows (100-150, 150-200, and 200-250 ms in Fig. 5A). Using specific latencies allowed us to more clearly see neural activity that was aligned to the upcoming saccade. As in Fig. 3, we compared scenarios when there were differing amounts of prior information based on position (angular position is near vs. opposite the PD), and also when the actual upcoming saccade was toward vs. away from neurons’ PDs. This allowed us to track how neural activity reflected both prior information and the selected upcoming saccade over time.

**Figure 5:**
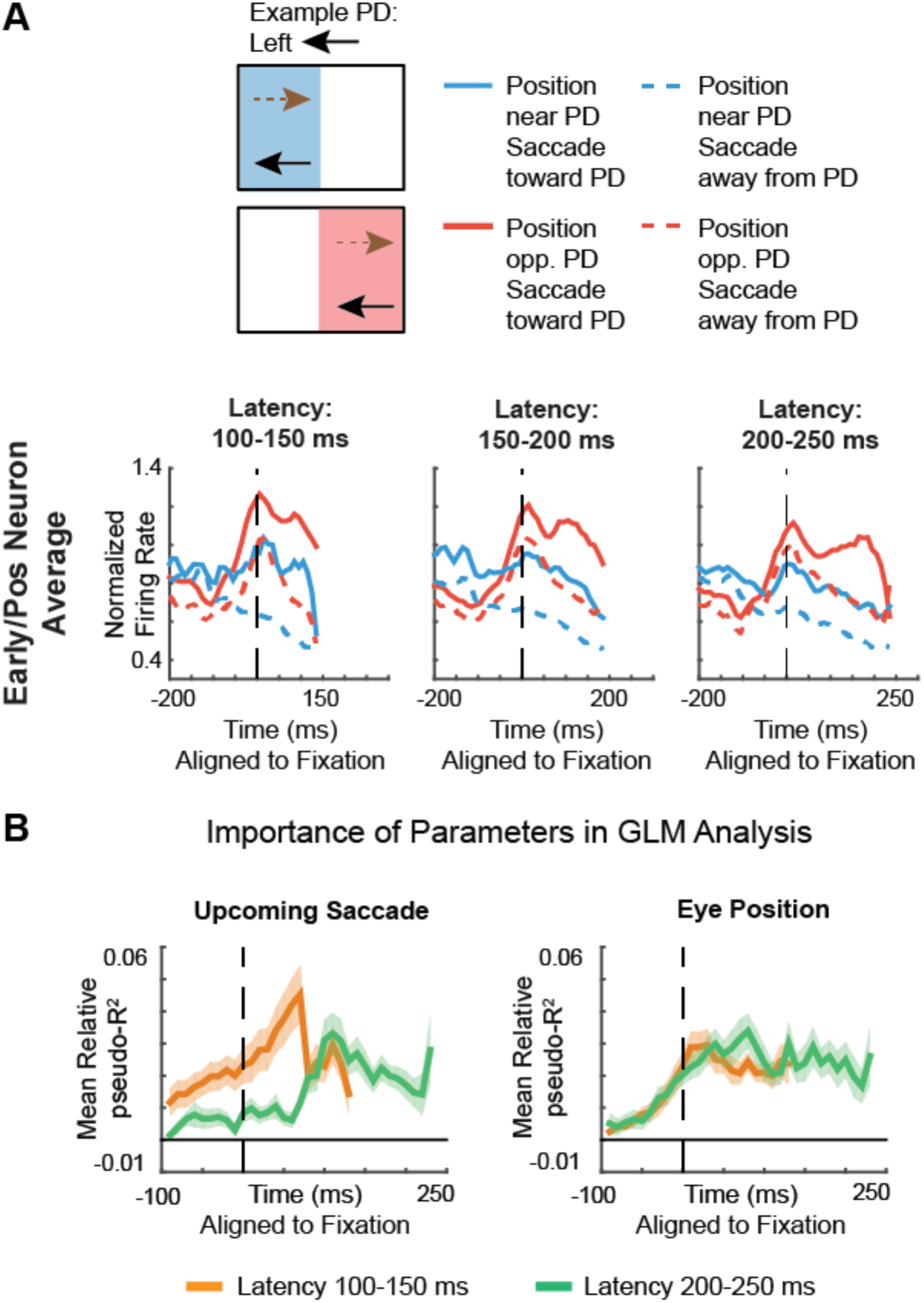
Evolution of activity over time. **(A)** PETHs, aligned to fixation onset, of normalized averaged activity of Early/Pos neurons. Blue lines are those with a starting angular eye position near the PD (unlikely that upcoming saccade will be toward PD). Red lines are those with a starting angular eye position opposite the PD (likely that upcoming saccade will be toward PD). Separate PETHs are constructed for saccades with latencies from 100-150 ms (left), 150-200 ms (middle), and 200-250 ms (right). **(B)** Importance of parameters in the generalized linear model, across time, aligned to fixation onset, for Early/Pos neurons. The mean relative pseudo-R^2^ (across Early/Pos neurons) of the upcoming saccade (left) and eye position (right) covariates are shown. We separately determine parameter importance for saccades with latencies of 100-150 ms (orange) and 200-250 ms (green). Shaded areas represent SEMs.

We first analyzed how activity reflected the actual upcoming saccade (solid versus dashed lines in Figs. 5A and S6A). Early on, around the time of fixation, there was already a differentiation of activity related to the actual saccade that would occur, when controlling for position. As time progressed towards saccade onset, this activity difference increased. Interestingly, two activity peaks were often visible when the upcoming saccade was into the PD – one peak around the time of fixation and another peak about 50 ms before the upcoming saccade onset.

We then analyzed how activity reflected position over time (red versus blue lines in Figs. 5A and S6A). As we saw before, there was differentiation of activity related to position early on. Interestingly, this differentiation of activity based on position continued until the upcoming saccade. That is, activity near the upcoming saccade reflected a mixture of prior information and information about the saccade that would actually occur.

To be more rigorous, we also used a generalized linear model (GLM) approach (Figs. 5B and S6B). This model can control for confounding factors in the PETH-based analyses, such as correlations with the previous saccade, and can better disentangle correlations between eye position and upcoming saccades. Additionally, in the GLM, the saccade and positions are no longer only characterized by angles, as they were in the PETHs. As input variables to the model, we included the eye position, upcoming saccade vector, upcoming saccade velocity, and previous saccade vector (see *Methods*). This model-based analysis allowed us to directly estimate the importance of saccade and position variables for predicting neural activity.

The GLM analysis confirmed our results from the PETHs. The actual upcoming saccade was already significantly predictive of neural activity at the time of fixation. The importance of the upcoming saccade parameter grew until about 50-100 ms before saccade onset. The importance of the position parameter increased until the time of fixation, and then stayed approximately constant until the time of saccade onset. That is, the prior continued to have an influence on activity throughout the entire saccade planning process. Combining the results from both parameters, while activity continuously relates to both prior information and the actual saccade, as time elapses, a greater proportion of the activity relates to the actual saccade that will occur.

### Relationship between early activity and the final selected saccade

Above, we observed that early activity around the time of fixation was predictive of the actual saccade that would occur, beyond the effects of position (Fig. 5). In order to better understand how early activity relates to the final saccade that is selected, we further examined a few scenarios.

First, we looked at how early neural activity related to the final saccade, depending on the latency of the saccade. PETHs showed that for longer latency (200-250 ms) saccades, there was only a small activity difference between saccades toward versus away from neurons’ PDs (when controlling for position; Figs. 5A and S6A). However, for shorter latency (100-150 ms) saccades, there was a larger activity difference based on whether the resulting saccade was into the PD. The GLM analysis confirmed these results. At the time of fixation, there was only a small amount of unique information about the upcoming saccade for longer latency saccades, while there was more information for shorter latency saccades (Figs. 5B and S6B). Thus, when the early neural activity contains more information about the upcoming saccade, it happens faster.

Next, we looked at the scenario when the next saccade is to the target. Behaviorally, we observed that when the next saccade is to a nearby target (meaning there was probably a strong “likelihood”; see *Behavior* subsection), the prior had less influence on behavior (Fig. 1G). We investigated the neural data to determine whether early neural activity was less informative about the upcoming saccade when there was a strong likelihood. We would expect that if the saccade decision is made based on visual processing that is to occur later, then early neural activity shouldn’t be predictive of the actual saccade. PETHs show that this is the case. When the saccade is to a nearby target (< 10° away), there is no longer an early activity difference based on whether the upcoming saccade is into the PD (when controlling for position; Figs. 6A and S7A). We find the same conclusion in a GLM-based analysis. When looking at all saccades, we can see that the actual upcoming saccade is predictive of neural activity, beyond the effects of position. However, when going to a nearby target, the actual upcoming saccade is not predictive of early neural activity (Figs. 6B and S7B). That is, the neural activity is not informative about the actual upcoming saccade beyond its information about position. Note that this result is not due to latency differences (Fig. 5), as saccades to the target have shorter latencies [12]. Thus, neural activity around the time of fixation does appear to act like a Bayesian prior about the upcoming saccade.

**Figure 6:**
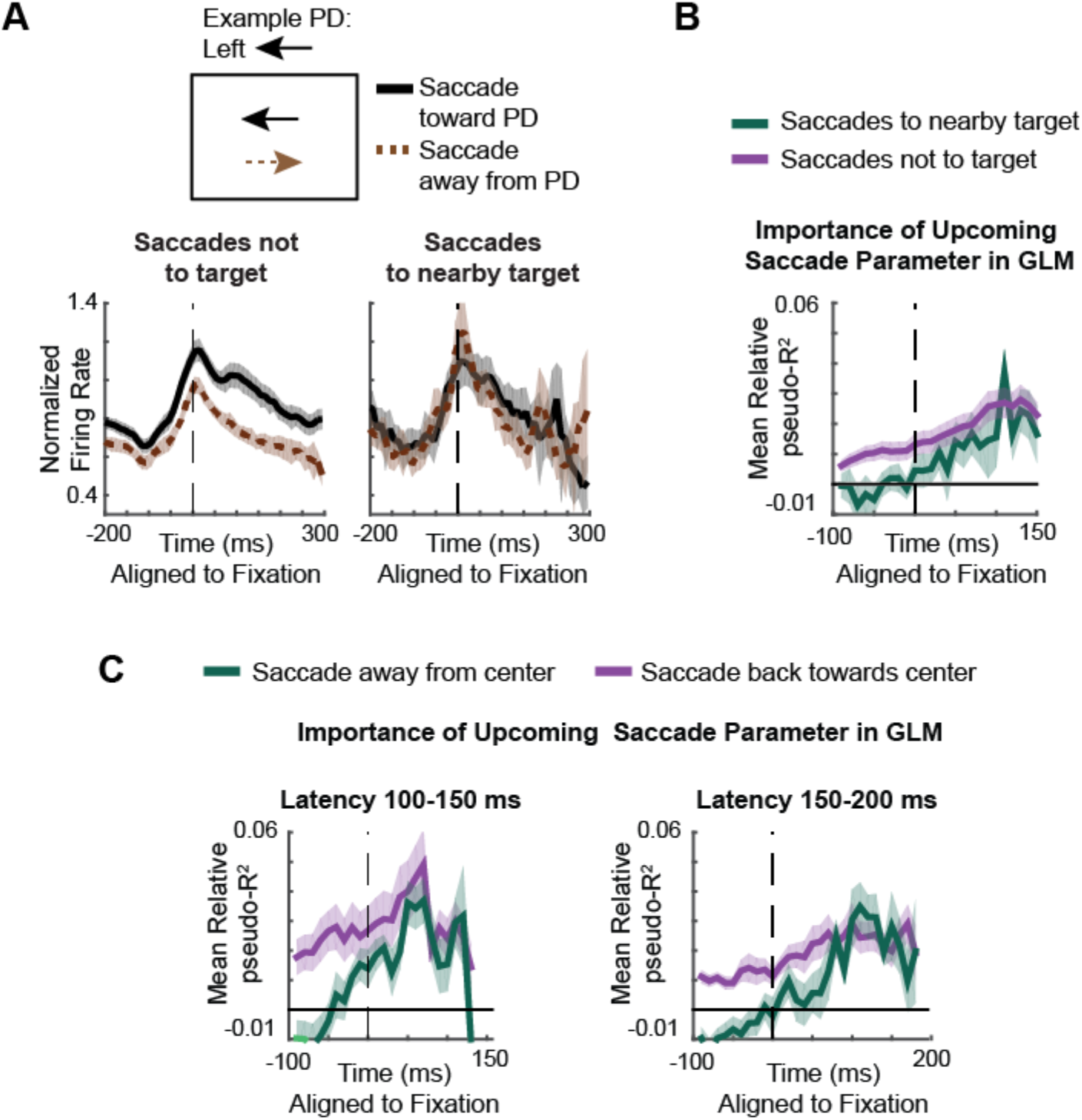
Relationship between early activity and the final selected saccade. **(A)** PETHs of Early/Pos neurons, aligned to fixation, of saccades toward the preferred direction (PD; black) versus away from the PD (brown, dashed). To control for position, we only use saccades starting from positions opposite the PD. On the left, we only include saccades not to the target. On the right, we only include saccades that go to a nearby target (< 10° away). **(B)** Importance of the upcoming saccade parameter in the generalized linear model, across time, aligned to fixation. The mean relative pseudo-R^2^ (across Early/Pos neurons) was determined for saccades not to the target (purple) and to a target < 10° away (gray). Note that the GLM results were only plot until +150ms, as the results got very noisy since there are limited saccades with latencies > 150 ms. **(C)** Importance of the upcoming saccade parameter in the generalized linear model, across time, aligned to fixation. The mean relative pseudo-R^2^ (across Early/Pos neurons) was determined for saccades back towards the center (purple) and saccades away from the center (gray). Shaded areas represent SEMs. Note that for this comparison, we cannot do a PETH analysis like in previous scenarios, because saccades back to the center, from an angular position opposite the PD, will always be toward the PD (we can’t compare saccades toward versus away from the PD while controlling for position).

Finally, we looked at scenarios when the upcoming saccade is in agreement with the prior information (back towards the center) versus disagreement with the prior information (further away from the center). It is likely that if the saccade ends up going against the prior, that the decision was likely to be primarily based on visual information gathered later (not the prior information). However, if the saccade is in agreement with the prior, then it is possible that this prior information was used in the saccade decision. We analyzed whether the early neural activity was informative about the final saccade decision using a GLM-based analysis. We controlled for latency given that saccades back towards the center have lower latency. Indeed, we found that neural activity preceding fixation was less informative about the upcoming saccade for saccades away from the center (against the prior; Figs. 6C and S7C). This again supports a Bayesian interpretation of the early activity representing a position-based prior; when the final saccade decision is not based on the prior, this early activity is not very informative about that saccade.

### “Late” Neurons

While the focus of this paper has been on Early neurons, there were also many “Late” neurons. Late neurons were defined as those significantly modulated by the upcoming saccade in a GLM analysis in the 100 ms before saccade onset, but that were not significantly modulated by the upcoming saccade around the time of fixation (i.e., they were not Early neurons). 51/180 (28%) of recorded neurons were classified as Late neurons. As seen from PETHs (Figs. 7A and S8) and our GLM analysis (Fig. 7D), these neurons’ activities start to differentiate based on the upcoming saccade after fixation onset. They also appear to be better aligned to saccade onset than fixation (Figs. 7A and S8). Thus, there is a separate group of neurons with a later time of saccade selectivity.

**Figure 7:**
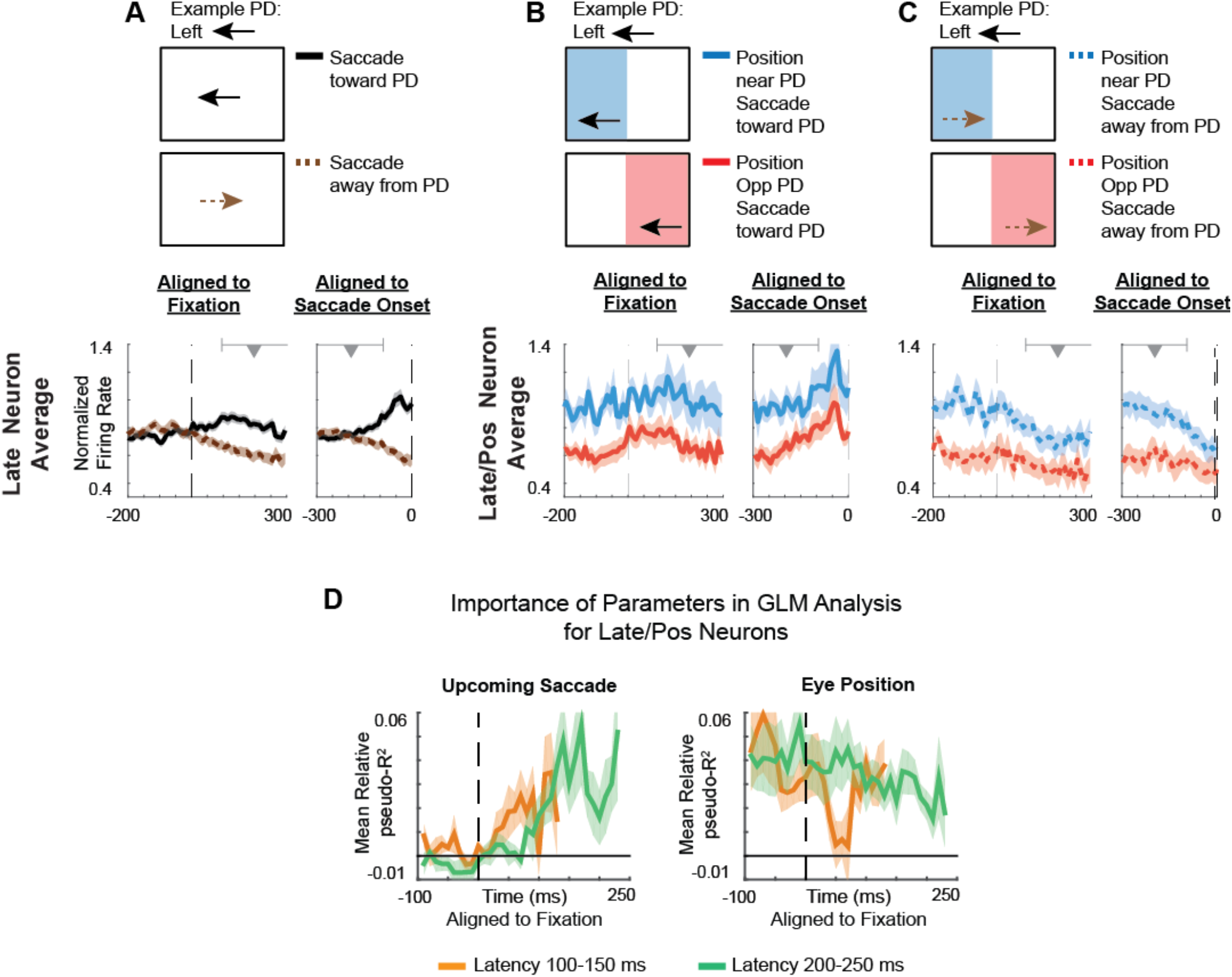
Late Neurons. **(A-C)** PETHs, aligned both to fixation (left part of each column) and the upcoming saccade onset (right part of each column). Ranges of saccade/fixation onset times are shown above the PETHs as in Fig. 2. **(A)** PETHs of saccades toward the preferred direction (PD; black) versus away from the PD (brown, dashed). PETHs are averaged across Late neurons. **(B)** PETHs of saccades toward the PD, with a starting angular eye position near the PD (unlikely that upcoming saccade will be toward PD; blue) versus an angular position opposite the PD (likely that upcoming saccade will be toward PD; red). For example, if the PD is to the left, angular positions near the PD will be on the left side of the screen (see *Methods* for details). PETHs are averaged across Late/Pos neurons (Late neurons that are also significant for position). **(C)** PETHs of saccades away from the PD, with a starting angular position near the PD (blue, dashed) versus an angular position opposite the PD (red, dashed). PETHs are averaged across Late/Pos neurons. **(D)** Importance of parameters in the generalized linear model, across time, aligned to fixation. The mean relative pseudo-R^2^ (across Late/Pos neurons) of the upcoming saccade (left) and eye position (right) covariates are shown. We separately determine parameter importance for saccades with latencies of 100-150 ms (orange) and 200-250 ms (green). Shaded areas represent SEMs.

Do these Late neurons also have prior information related to position? As we did previously for Early neurons, we used a GLM to identify neurons that were significant for position around the time of fixation; these would be candidates for having prior information based on position. We found that 13/51 (25%) of Late neurons were significant for position, a much lower percentage than for Early neurons. Moreover, when we further investigate these “Late/Pos” neurons, it is clear that they are not representing prior information about the upcoming saccade in the manner of Early/Pos neurons. For instance, the importance of the position parameter stays relatively constant over time (Fig. 7D; note that the transient decrease in the orange trace is primarily driven by outliers). More importantly, PETHs show that average activity is actually higher when the angular position is near neurons’ PDs (blue trace), which is when the upcoming saccade is less likely to be in the PD (Fig. 7B,C). Thus, separate from Early neurons, there is another subset of neurons that only represents saccades after fixation onset and doesn’t represent prior information based on eye position.

## Discussion

Here, during a self-guided search task we found a subset of (“Early”) neurons that represented prior information about the upcoming saccade, often before the onset of fixation. This early neural activity related to prior information was predictive of the final selected saccade in a Bayesian manner. As time elapsed towards the upcoming saccade, prior information continued to have an influence on these neurons’ activities, but the activities evolved so that activity became more related to the final saccade selection. In a separate subset of (“Late”) neurons, activity only related to the final selected saccade and not prior information. Our findings demonstrate how prior information influences and evolves into definitive saccade plans.

Our findings have some overlap with the results of Phillips and Segraves [4], who also studied FEF during a natural scene search task. Like us, they found early saccade predictive activity in many neurons, sometimes prior to fixation. In their study, they also found that many neurons’ activities were predictive of future saccades (not just the upcoming saccade), which they called “advanced predictive activity”. When they split neurons into two subsets depending whether the neurons had advanced predictive activity or not, they found that neurons with advanced predictive activity also became selective for the upcoming saccade significantly earlier. Their separation based on whether neurons had advanced predictive activity (i.e. whether their activity predicted future saccades), may thus overlap with our separation into Early and Late neurons.

Our findings suggest a link between previous studies showing pre-target preparatory activity in constrained tasks and studies showing advanced saccade planning during self-guided saccades. Past studies have shown that superior colliculus neurons had higher activity prior to target onset when there was a higher probability the target would be shown in the neurons’ PDs [5,6]. This parallels our finding that Early/Pos neurons had higher activity when there was a greater probability of the upcoming saccade being in their PDs. Additionally, during self-guided search, researchers have provided evidence for FEF planning more than one saccade in advance [4,10]. These advanced plans could be reflected by Early neurons, which are predictive of the upcoming saccade before gathering new information. Thus, there may be a common mechanism, where Early neurons are involved in preliminary planning, whether based on saccade probabilities or some saccade sequence planned in advance.

### Visual remapping

A potential alternative explanation for the early activity is visual remapping [23–25]. Rather than being related to the likely upcoming movement, this early activity could be related to the visual scene that is about to be brought into the neurons’ receptive fields. For example, let’s say a neuron has a receptive field to the left. When the monkey is looking on the right side of the screen, there may be more interesting features in the neuron’s receptive field (to the left), as compared with when the monkey is looking on the left side of the screen, when the receptive field may include some visual space outside (to the left) of the scene.

However, there are a few reasons we believe visual remapping is unlikely to explain our results. First of all, in natural scene search, we have previously found that neural activity related to movement dominates that related to visual features. More specifically, we have found no evidence that FEF activity is modulated by visual features that are salient or task-relevant, beyond these features’ correlations with upcoming movements [12,13,26]. Previous demonstrations of visual remapping effects have been elicited by flashing a salient stimulus against a uniform background [23–25], which did not occur in our experiment. Moreover, if the early activity were due to visual remapping, we would expect activity differences between when the target is and isn’t being brought into the RF. However, we do not see an activity difference in this scenario (Fig. 6A). Finally, visual remapping starts to occur prior to the previous saccade onset [27,28], while our “early planning” signal typically happens around the time of the previous saccade onset (Fig. S9). Thus, it is unlikely that the early activity is primarily a visual remapping signal.

### Neural activity related to eye position

Here, we assumed that the neural activity of Early neurons related to eye position was used for preliminary planning. This is in line with previous research showing that a lower stimulation threshold in FEF was required to elicit saccades opposite of the current eye position [29], which suggested that eye position biases upcoming saccades. However, it is possible that FEF activity related to position was used for computations other than, or in addition to, saccade planning. For instance, Cassanello and Ferrera [30] found that there was generally greater activity in FEF neurons when the initial eye position was opposite the neurons’ PDs. However, they argued that this position-based modulation of activity could allow vector subtraction, with the purpose of keeping a memory of the target location across saccades. It is important to note that a position signal could ultimately be used for multiple purposes. For instance, there could be multiple readouts of this position signal, one that is used for saccade planning, and another that does vector subtraction for the purpose of stability across saccades. Ultimately, given that Early/Pos neurons’ activities with respect to position matched the statistics of upcoming saccades, and given that FEF has a known role in saccade planning [1,31,32], it is improbable that these neurons were representing position solely for a purpose other than making saccade decisions.

Previous studies have also suggested that neural activity in superior colliculus (SC) is modulated by eye position in order to bias upcoming saccades. Pare and Munoz [11] found that burst neurons in SC had higher firing rates when the eye position was opposite the neurons’ PDs, as we found here for FEF. However, other studies in SC [33,34] found the reverse result (although in different tasks) – that firing rates were generally higher when the eye position was in the same direction as the neurons’ PDs. It is thus possible that a subset of SC neurons use position for preliminary planning. Given the effect of eye position on saccade latencies (Fig. 1), it makes sense that it would affect neural activity related to saccade planning throughout the oculomotor system.

How does the FEF have access to eye position information to use for saccade planning? Given that Early/Pos neurons are modulated by the fixation position prior to the start of fixation, these neurons cannot be using a sensory eye position signal. Rather, we suspect that this information is computed based on a corollary discharge signal of the saccade plan. A strong candidate source for this signal is from the superior colliculus via mediodorsal thalamus [35,36]. In fact, when the collicular thalamo-cortical pathway is blocked, monkeys are not able to successfully make sequences of saccades [35,36]. Thus, we would hypothesize that blocking this corollary discharge pathway would interfere with Early/Pos neurons’ representation of prior information.

Additionally, some Late neurons also had activity that was modulated by position, although not in a manner that appeared to be related to the upcoming saccade. It is possible that the source for this position information could be a proprioceptive signal [37], possibly derived from eye position information in area 3a of the somatosensory cortex, and connections from area 3a to FEF [38].

### Cell types

While we split neurons into Early and Late neurons, these neurons may lie along a spectrum rather than being discrete classes. While the majority of Early neurons had activity predictive of the upcoming saccade both near the time of fixation and near the time of the saccade (e.g. top row of Fig. 2), some Early neurons did not have much saccade-predictive activity near the time of the upcoming saccade (e.g. third row of Fig. 2). Late neurons only had saccade-predictive activity near the time of the upcoming saccade. Thus, it could be the case that neurons lie on a spectrum from having saccade-predictive activity only around fixation to only around the upcoming saccade, with many neurons having a mixture (Fig. S10).

Classically, researchers have used a memory-guided saccade task to categorize FEF neurons as having visual, delay, and/or movement activity [39,40] (although see [41] for recent work revisiting these classifications). As past work has shown sensory to motor transformations from visual to movement cells [42], it is interesting to speculate how Early and Late neurons relate to these classical cell types. One simple explanation could be that these neurons lie on the classical visual to movement spectrum depending on how much of their saccade-predictive activity is near the time of fixation versus near the time of saccade. However, it is probably not that simple. Phillips and Segraves [4] found that similar proportions of (classically defined) visual and visuomovement cells had advanced predictive activity. Moreover, the majority of our Late neurons had activity modulated by visual scene onset (Fig. S11), suggesting that Late neurons are not purely movement related. Thus, it does not appear that Early and Late neurons cleanly map onto classical FEF cell types.

### Probability distributions

When averaging activity across saccades, the activity of Early/Pos neurons was similar to the full continuous probability distribution of upcoming saccades (Fig. 4C,D). However, it did not match exactly. The distribution of neural activity did not have the peaks at around 135° and 225° like the behavioral distribution. This could be because the activity of these FEF neurons only serves as general prior information to direct saccades towards the center. In this scenario, other oculomotor structures could relate to other aspects of the prior. Alternatively, we only recorded a limited number (28) of Early/Pos neurons, and it is possible that recording a large number would reveal a distribution that more precisely matches behavior. Further experiments while recording even more neurons will be necessary to differentiate these possibilities.

Demonstrating that neural activity relates to a full, continuous probability distribution of saccade directions extends previous work showing that neural activity in the oculomotor system reflects the probabilities of upcoming saccades when deciding between a small number of discrete targets [5–7,43,44]. Importantly, because our results were based on averaging across saccades, we do not know whether the FEF population, prior to single saccades, reflects the probability distribution of the upcoming saccade. An alternative explanation is that the population always makes preliminary plans for a single saccade, and when averaged across saccades, these individual plans create a distribution. In the future, it would beneficial to simultaneously record many FEF neurons and perform a single-trial decoding analysis (as in [45]) to determine whether probability distributions or individual plans are represented prior to single saccades.

### Bayesian decision-making

There is a large literature demonstrating that decision-making for movement is approximately Bayesian [46,47]. Moreover, it was recently shown that the smooth pursuit region of FEF (FEF_SEM_) has activity that maps onto approximately Bayesian behavior [48]. Here we showed that neural activity relates to aspects of this Bayesian decision-making process during self-guided saccades. The early activity of Early/Pos neurons (which reflected prior information) was less predictive of the final saccade decision when there was stronger visual (“likelihood”) information (when the next saccade was to a target). Moreover, it is not only the early activity (around the time of fixation) that may relate to Bayesian decision-making; we can also view the evolution of Early/Pos neurons’ activities through a Bayesian framework. Given that the activity of Early/Pos neurons is continuously related to prior information, and increasingly related to the final decision over time, the activity may represent a posterior distribution. That is, it could represent the current beliefs about the upcoming saccade, based on the prior information and visual likelihood information. It would be valuable to do future experiments with more controlled visual information, to further understand FEF’s role in Bayesian computations for saccade decisions.

Additionally, it would be interesting to understand how this posterior gets transformed into a selected plan, which is reflected in the activity of Late neurons. From a computational perspective, Kim and Basso [49] showed that in SC, a Bayesian maximum a posteriori (MAP) model better predicted selected saccades than winner-take-all or population vector average models. From a neurobiological perspective, the neural circuits involved in transforming prior information to selected saccades remain unclear. One possibility is that Early neurons project to Late neurons within FEF to influence the saccade plan. Another possibility is that Early neurons project to neurons in SC, which then go on to influence the saccade plan. Both possibilities could also happen simultaneously. Clearly, our work could be explained by a large number of different circuit models. Future work should aim to elucidate the circuit mechanisms behind the transformation from prior information to definitive saccade plans.

## Methods

Many of the methods here, especially for neural data analysis, are the same as in our other recent manuscripts [12,45], and are described in the same way.

### Behavioral Paradigms

#### Experiment

Two monkeys (Monkeys J and K; in previous papers referred to as M15 and M16 [12,13]) freely searched for an embedded Gabor target in a natural scene background, as in [12,13]. They were rewarded for fixating near the target for 200 ms. If they did not find the target after 20 saccades, the trial ended.

#### Eye tracking

Eye movements were tracked with an infrared eye tracker (ISCAN Inc., Woburn, MA, http://www.iscaninc.com/) at 60 Hz.

#### Saccade detection

The start of saccades was determined by when the velocity of eye movements went above 80 degrees / sec. The end of saccades was marked by when the velocity fell below 100 degrees / sec. Saccades could only be detected after an intersaccadic interval (latency) of 90 ms. To be conservative about saccades, we only included saccades of at least 5 degrees (so that noise in the eye tracker was not classified as a saccade). Saccades longer than 80 degrees or with duration longer than 150 ms were discarded as eye-blinks or other artifacts.

### Neural Data Acquisition and Preprocessing

Monkeys J and K were implanted with a 32 channel chronic electrode array (Gray Matter Research, Bozeman, MT, USA) over the frontal eye field (FEF). The depth of each individual tungsten electrode (Alpha-Omega, Alpharetta, GA) could be independently adjusted over a range of 20 mm. Details about recording locations can be found in [12]. While discussing recording locations, we want to briefly note that there were many instances in which both Early and Late neurons (see *Results*) were recorded from the same electrodes at the same depths.

Automatic spike sorting with some manual correction was performed offline using the Plexon Offline Sorter (Plexon, Inc., Dallas, TX, USA). Because any given electrode was often left in place for multiple days, we often recorded from the same neuron across sessions. To make use of this, we combined data from units that persisted across recording sessions on different days. To do this, we manually compared spike waveforms from units recorded at the same site on different days. Generally, we merged units sharing waveform shape (rise/fall characteristics, concavity/convexity, etc.), and time course. Ambiguous cases were not combined.

In Monkey K, we stimulated with the electrodes to verify FEF location (details in [12]). Monkey J continues to be used in ongoing experiments. Since microstimulation would lower the impedance of the array electrodes and harm our ability to record neurons, we have not yet administered microstimulation in this animal. While we were not able to confirm the electrode placement for Monkey J, the array was in the same stereotactic location, and the response patterns were very similar. Thus (in addition to having the array stereotaxically above FEF), we used functional measures to include neurons that were likely in FEF (as in [13]). We only included neurons that either had visual onset activity or movement activity. To determine whether there was visual onset activity, we compared neural activity in the 100 ms prior to image onset with activity 50 to 150 ms after image onset, to see whether there was a significant difference (Wilcoxon rank-sum test; p<0.005). To determine whether there was movement activity, we looked at peri-saccadic time histograms aligned to the start of the upcoming saccade, binned into 8 angular directions (according to saccade direction), with each bin subtending 45 degrees. In any bin, we tested whether there was a significant difference between activity in the 100 ms around saccade onset and a baseline period 300-200 ms before saccade onset (Wilcoxon rank-sum test; p<0.005). In sum, while most of the neurons were likely in FEF, it is possible that some neurons were in nearby areas. After following the above inclusion criteria, we had 104 neurons from Monkey J and 76 neurons from Monkey K.

### Behavioral Analysis

We excluded saccades that started or ended outside of the boundaries of the screen. Behavioral data was combined across all sessions.

#### Statistics of movement

We defined the angular position, *ϕ_P_*, as the initial fixation location (prior to a saccade) relative to the center of the screen (Fig. 1c). We defined *ϕ* as the angular difference between the upcoming saccade direction, *ϕ_S_*, and the angular position (Fig. 1). That is, *ϕ* = *ϕ_S_* − *ϕ_P_*.

#### Latency effects

Latency was defined as the time from start of fixation to saccade onset. Latencies greater than 400 ms were excluded as outliers, as latencies of this duration could have been due to an undetected saccade.

We computed the mean latency of movements as a function of *ϕ*. When claiming that latencies were lower when making saccades opposite the angular eye position (when *ϕ* is close to 180°), we did the following test: We calculated the Pearson’s correlation between latency and |*ϕ* − 180° |. We then calculated the p-value associated with the correlation (using a 2-sided one-sample t-test).

We also analyzed differences in latencies between saccades that returned towards the center (|Φ-180°|<60°) and saccades away from the center (|Φ-180°|>120°), based on the distance of the eye position from the borders of the screen. To test whether the latency difference between towards-center and away-from-center saccades depended on the distance from the border, we used linear regression to fit the latency of saccades as a function of distance from the center. We then did a 2-sided unpaired t-test with unequal sample variances to analyze whether the slope was less (more negative) for towards-center saccades compared to away-from-center saccades.

### Neural Data Analysis

As in our behavioral analyses, we only included saccades that remained on the screen. Additionally, except when noted otherwise, we excluded the first saccade of each trial to remove the confound of the large visual onset driven by the appearance of the image.

#### Smoothed maps of neural activity

For many aspects of the following neural data analysis, we computed smoothed maps of neural activity in relation to some variable (position, previous movement, or the upcoming movement). For instance, we created a map of how neural activity varied over all positions on the screen, and a map of how neural activity varied in response to all upcoming saccade vectors. For our maps, we estimated the average firing rate at each point in space using weighted k-nearest neighbor smoothing. As an example, for the saccade variable (previous or upcoming), for each saccade we found the *k* nearest saccade vectors (based on Euclidean distance). We then averaged the firing rates associated with each of the *k* saccades, but with each weighted proportional to its distance from the given saccade to the *d* power.

The parameters (*k* and *d*) we used to generate the smoothed maps of neural activity are as follows. For the smoothed maps used in the generalized linear models (see section below), *k* = the smaller of 30% of the data points and 500, *d*=0. These parameters were found using cross-validation on held out data sets, in order to not inflate the number of significant neurons in the GLM analysis. For all other times, *k* = the smaller of 30% of the data points and 400, *d*=−0.5. These parameters were found using cross-validation on the current data sets in order to create as accurate maps as possible. Importantly, all results were robust to a wide range of smoothing parameters.

For any single variable (e.g. position) we can get the associated estimated firing rate, *θ_P_*, by looking up the firing rate for that position on the smoothed map. If, for instance, we want to get the estimated firing rates due to position in a time interval before every saccade, we would get a vector ***θ_P_***, which contains the estimate before each saccade. The same can be done to estimate the firing rates due to the upcoming saccades, ***θ_US_***, or previous saccades, ***θ_PS_***.

#### Determining Preferred Directions of Neurons

When determining the PD, we used the 100 ms preceding saccade initiation. Let ***Y*** be the vector of firing rates in that interval for every saccade. We fit a von Mises function to relate the movement directions to the firing rate due to movement:

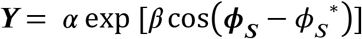

where ***ϕ_S_*** is the vector of upcoming saccade directions, and *α, β*, and *ϕ_S_*^*^ are the parameters that get fit. *ϕ_S_*^*^ is the PD of the neuron.

When estimating the PD, we only used the time period prior to the first saccade of each trial, when the eye position was in the center and there was not a previous saccade that was just ending, so these were not confounding factors. Moreover, this makes it so saccades used for estimating the PD were not included in the actual data analyses.

#### PETHs

When plotting PETHs of individual neurons, we plotted the mean firing rate across saccades. The error bars on PETHs are the standard error of the mean (SEM) across saccades. When plotting the PETHs averaged across neurons, we first calculated the mean firing rate (across saccades of the given condition) over time for each neuron. We then normalized this activity trace for each neuron by dividing by the maximum firing rate of the average trace (across all conditions). We then show the average of these normalized firing rates across neurons. Error bars are the SEM across neurons. The traces in Figs. 2, 3, and 7 (and corresponding supplemental figures) are smoothed using a 10 ms sliding window. The traces in Figs. 5 and 6 (and corresponding supplemental figures) are smoothed using a 30 ms sliding window, as there are fewer saccades in the conditions in those PETHs. For the PETHs aligned to fixation, only data obtained before the onset of the saccade are included.

PETHs were made for different categories of movements. For the PETHs, saccades toward the PD were defined as those that were within 60° of the PD. Saccades away from the PD were defined as those greater than 120° from the PD. Positions opposite the PD were defined as angular positions greater than 120° away from the PD. Positions near the PD were defined as angular positions less than 60° away from the PD.

#### Generalized Linear Model

To determine which variables were reflected in the neural activity, we used a Poisson Generalized Linear model (GLM). Let ***Y*** be a vector containing the number of spikes in the time interval we are considering, for every saccade. It has size *m x* 1. We aimed to predict ***Y*** based on several factors. Unless otherwise noted, we used the eye position, the previous saccade vector, the upcoming saccade vector, the peak velocity of the upcoming saccade, and a baseline term. More specifically, the covariate matrix ***X*** was:

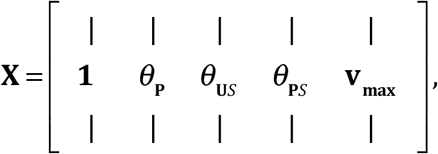

where ***θ_P_, θ_US_***, and, ***θ_PS_*** are generated from the smoothed maps (see *Smoothed maps of neural activity* above). Essentially, these covariates are the expected firing rates from position, upcoming saccade, and previous saccade (respectively) by themselves. Note that these covariates are not just based on angle, as they were when making PETHs – saccade amplitudes and the distance of the position from the center matter. **v_max_** is the vector of peak velocities of movements. The peak velocity was relative to the expected velocity given the main sequence [50], to control for the changes of velocity with saccade amplitude (as in [12]). When we run GLMs during different time intervals, we make separate smoothed maps for these time intervals. Note that when determining whether neurons were “Early” or “Late” (and Fig. S10), we only used the previous and upcoming saccade vectors (and a baseline term) as covariates.

Overall, the model that generates the firing rate (*λ*; also known as the conditional intensity function) can be written as:

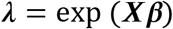

where ***β*** is a vector of the weights for each covariate that we fit, and ***X*** is the matrix of covariates, which is *z*-scored before fitting. If there are *j* covariates, then ***β*** has size *j x* 1. ***X*** has size *m x j*. Note the use of an exponential nonlinearity to ensure that firing rates are positive. The model assumes that the number of spikes, ***Y***, is generated from the firing rate, *λ*, according to a Poisson distribution.

We fit the model weights to the data using maximum likelihood estimation. That is, we found ***β*** that was most likely to produce the true spike output (assuming spikes were generated from the firing rate in a Poisson nature). Critically, we used (5-fold) cross-validation, meaning that the model was fit to the data using one set of data (the training set), and model fits were tested with an independent set of data (the testing set). Similarly, when calculating the test set covariates for saccade and position (described in *Smoothed maps of neural activity*), we only used k-nearest neighbors from the training set, to avoid overfitting.

To test whether an individual covariate significantly influenced neural activity, we first made sure that a simplified model with only that individual covariate had significant predictive power. To determine the value of a model fit, we used pseudo-R^2^ [51,52], a generalization of R^2^ for non-Gaussian variables, which is defined as:

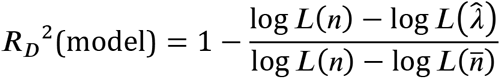

where log *L*(*n*) is the log likelihood of the saturated model (i.e., one that perfectly predicts the number of spikes), log 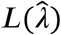 is the log likelihood of the model being evaluated, and log 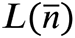 is the log likelihood of a model that uses only the average firing rate.

Then, in order to determine the importance of that covariate to the full model, we tested whether the full model predicts neural activity significantly better than a model where that covariate is left out (reduced model). To compare the fits between the reduced model (model 1) and full model (model 2), we used relative pseudo-R^2^, which is defined as:

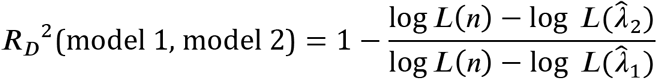

where log 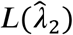 is the log likelihood of the full model and log 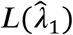 is the log likelihood of the reduced model.

To determine significance, we bootstrapped the fits to create 95% confidence intervals, and checked whether the lower bounds of these confidence intervals were greater than 0. Note that the pseudo-R^2^ and relative pseudo-R^2^ values can be less than 0 due to overfitting.

#### Population activity over time averaged across trials

For each neuron (of the category we were plotting), we calculated the firing rate as a function of the relative angular position (Fig. 4). We defined the relative angular position as the difference between a neuron’s PD and the eye position (the PD minus the angular eye position). We then normalized each neuron by dividing by its mean firing rate, and then averaged the normalized activity across neurons. We then smoothed the activity for plotting using the parameters from the smoothed maps. As with the PETHs, only data obtained before the onset of the upcoming saccade are included.

We also made variants of the above plot. We made plots in which only saccades near or far from the border (split based on median distance to the nearest border) were included. To control for the correlation between the previous and upcoming saccades, we made a plot where saccades were only used if the angle between previous and upcoming saccades was less than 90°.

#### Visual Activity

We determined whether cells had “visual” activity based on the response when the visual scene was first displayed. For this analysis, for each neuron, we only included trials in which the first saccade after scene onset was away from the neuron’s PD, to remove the potential confound of upcoming movement activity. Neurons were classified as “visual” if they met two criteria. First, activity from 50-150 ms after scene onset had to be significantly different (*p* < .05 using a Wilcoxon Rank Sum test) than activity in the 100 ms before scene onset. Second, to ensure a relatively sharp rise in activity after scene onset, neurons’ activities needed to increase by at least 35% within a 50 ms window (e.g. from 40-90 ms after scene onset, or 50-100 ms).

## Acknowledgements

We would like to thank Sara Caddigan and Emily Berthiaume for technical assistance. We would like to thank Hugo Fernandes, Pavan Ramkumar, Matt Perich, and Lee Miller for helpful discussions and comments. For funding, we would like to thank NIH F31 EY025532 and NIH R01 EY021579.

## Supplementary Figures

**Figure S1:**
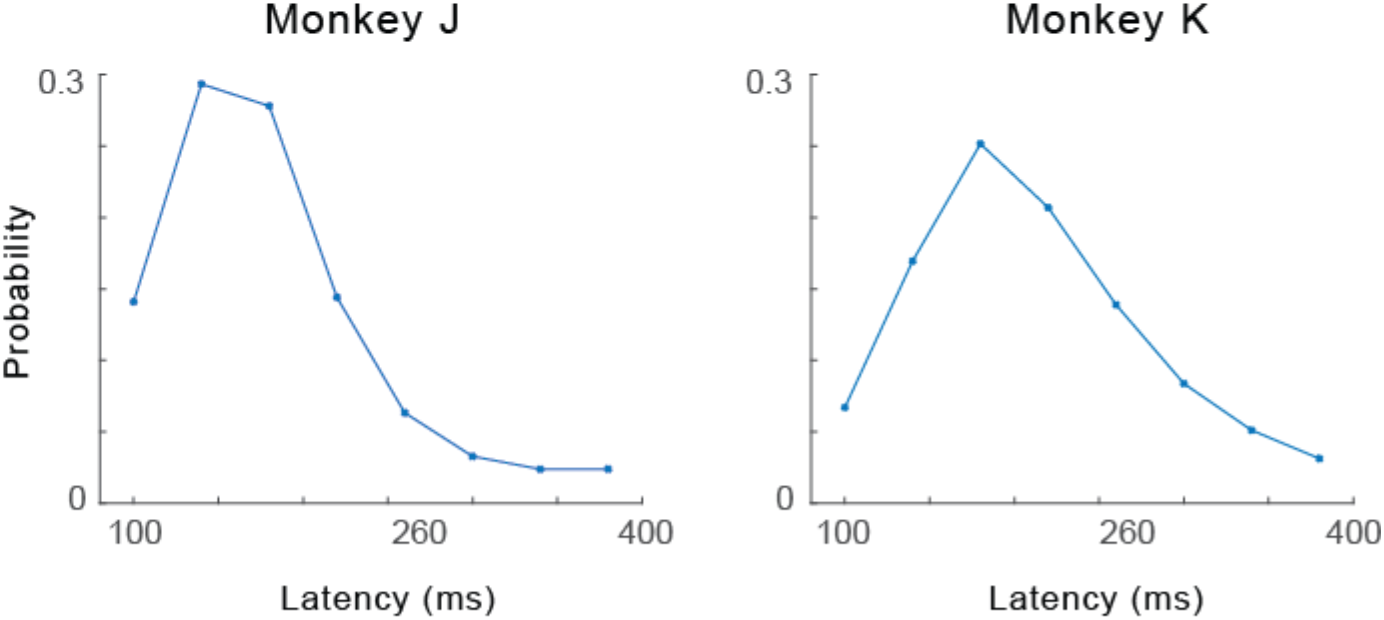
Distribution of saccade latencies. For each monkey, we show the distribution of saccade latencies (intersaccadic intervals). Probabilities were calculated within 40 ms bins from 80-400 ms, so the data point at 260 ms represents the probability of latencies from 240-280 ms.

**Figure S2:**
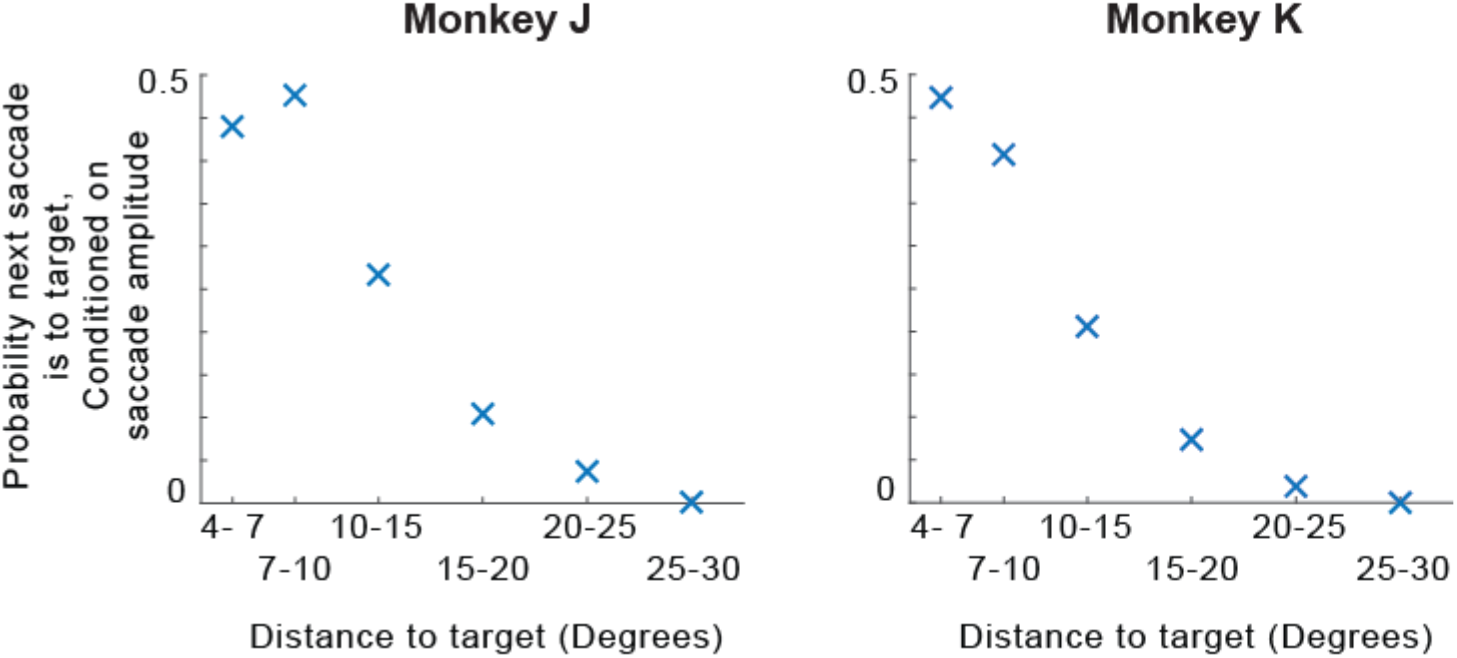
Probabilities of saccades to the target. For each monkey, when the monkey is at different distances to the target (x-axis), we show the probability that the next saccade is to the target, while controlling for saccade amplitude. To control for differing probabilities of saccade amplitudes, we divide by the probability of making a saccade of the amplitude required to acquire the target, regardless of whether the saccade is in the direction of the target. Thus, the y-axis can be viewed as the probability of making a saccade to the target, relative to the probability of making any saccade of similar amplitude.

**Figure S3:**
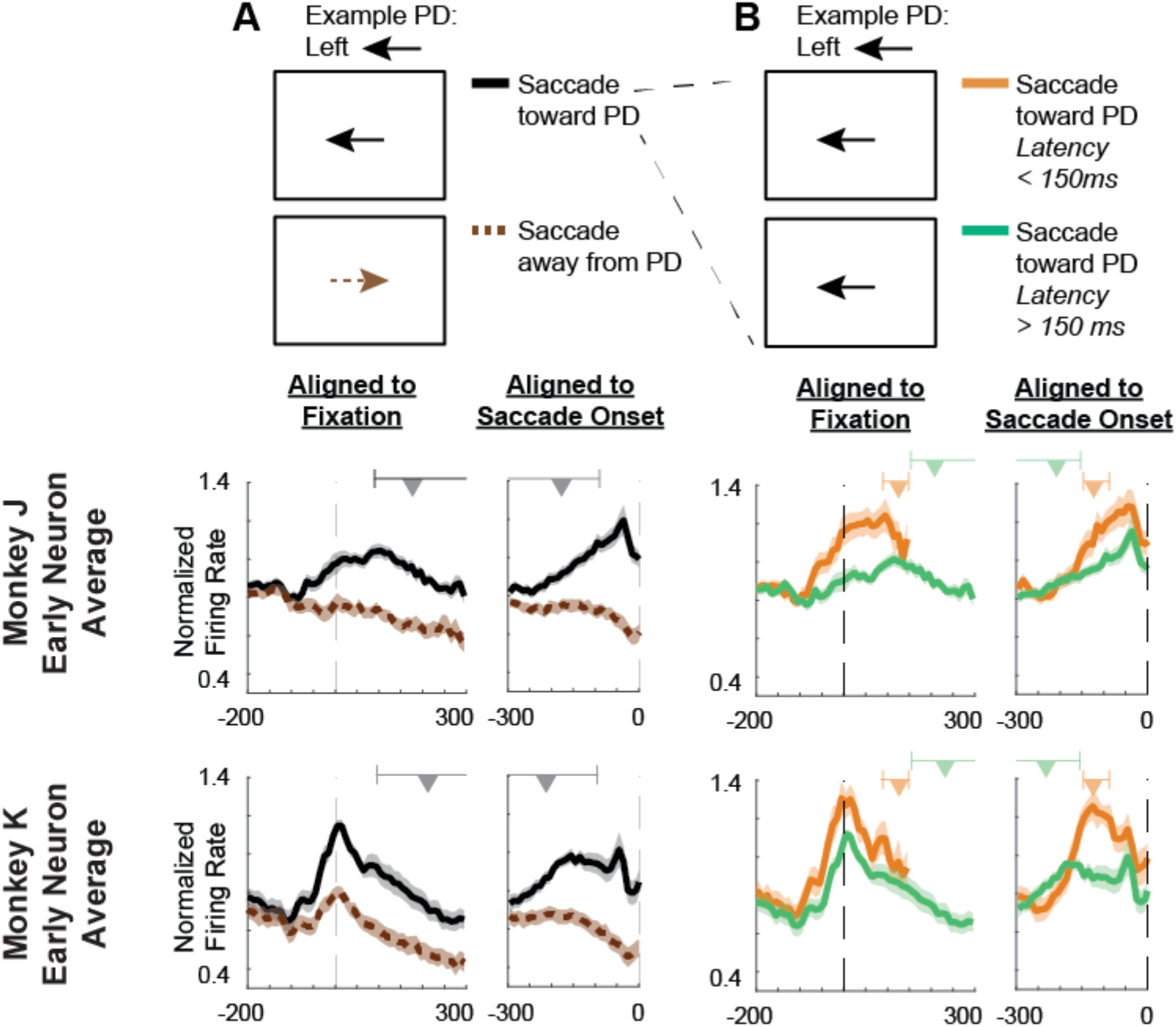
Early times of saccade selectivity, and neural differences related to saccade latencies, for individual monkeys. Peri-event time histograms (PETHs), aligned both to fixation (left part of each column) and the upcoming saccade onset (right part of each column). **Top Row of PETHs**: Normalized averages of Early neurons from Monkey J. **Bottom Row**: Normalized averages of Early neurons from Monkey K. **(A)** PETHs of saccades toward the preferred direction (PD; black, solid) versus away from the PD (brown, dashed). Above the PETHs aligned to fixation, we show the range of 95% of saccade initiation times (the upper end of this range is larger than the x-axis limit). Above the PETHs aligned to saccade onset, we show the range of 95% of fixation onset times (the lower end of the range is below the x-axis limit). The triangles represent the median times. **(B)** PETHs of saccades toward the PD (like the black trace in panel A), divided further based on saccade latency. Saccades with latencies less than 150 ms are shown in orange while saccades with latencies greater than 150 ms are in green. Above the PETHs, ranges of fixation/saccade times are shown separately for the separate traces. In all figures, for the plots aligned to fixation, only data obtained before the onset of the saccade are included in the PETHs.

**Figure S4:**
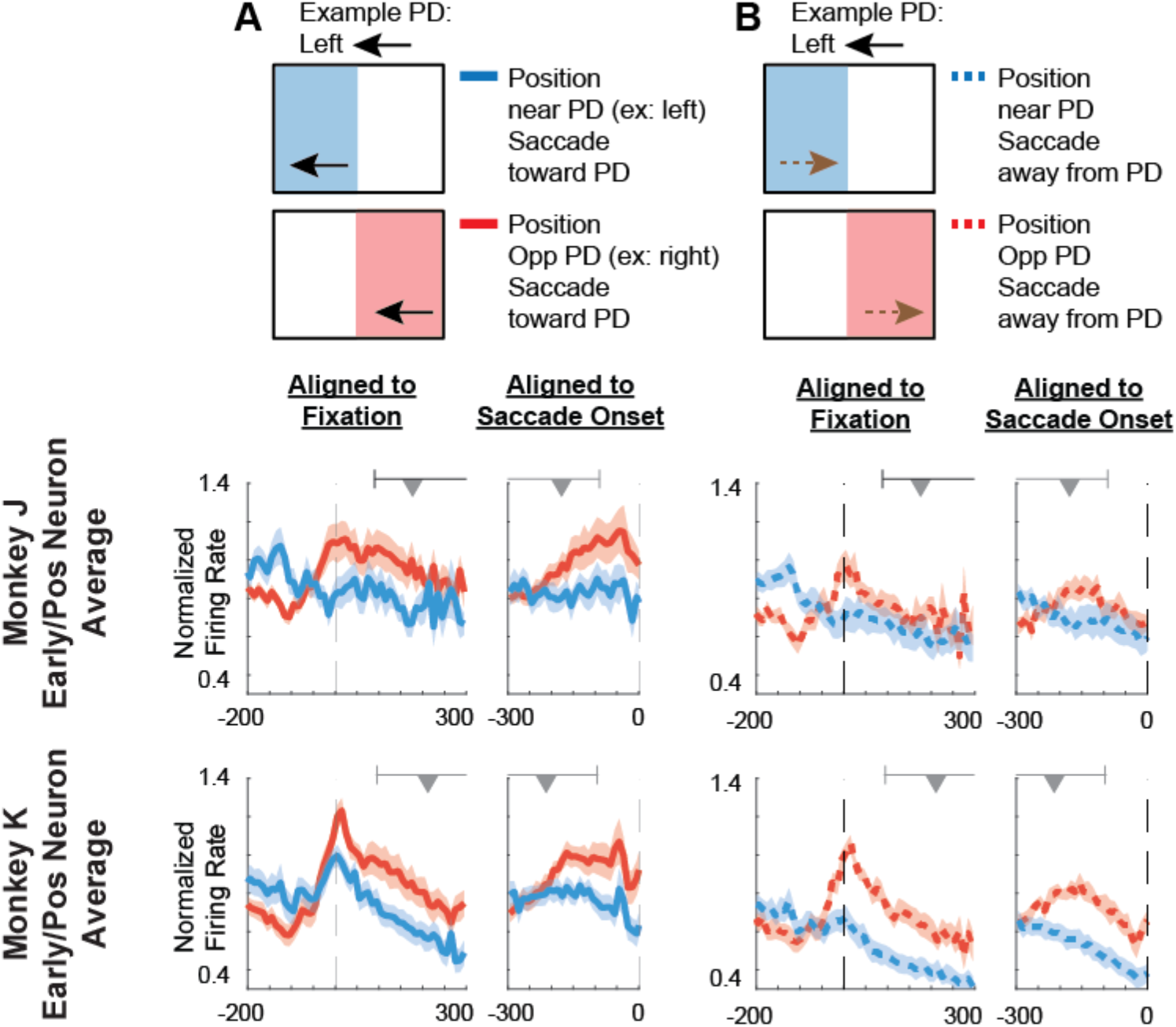
Increased activity in positions that are more likely to result in saccades toward the PD, for individual monkeys. Peri-event time histograms (PETHs), aligned both to fixation (left part of each column) and the upcoming saccade onset (right part of each column). Ranges of saccade/fixation onset times are shown above the PETHs as in Fig. 2. **Top Row of PETHs**: Normalized averages of Early/Pos neurons from Monkey J. **Bottom Row**: Normalized averages of Early/Pos neurons from Monkey K. **(A)** PETHs of saccades toward the PD, with a starting angular eye position near the PD (unlikely that upcoming saccade will be toward PD; blue) versus an angular position opposite the PD (likely that upcoming saccade will be toward PD; red). For example, if the PD is to the left, positions near the PD will be on the left side of the screen (see *Methods* for details). **(B)** PETHs of saccades away from the PD, with a starting angular position near the PD (blue, dashed) versus an angular position opposite the PD (red, dashed).

**Figure S5:**
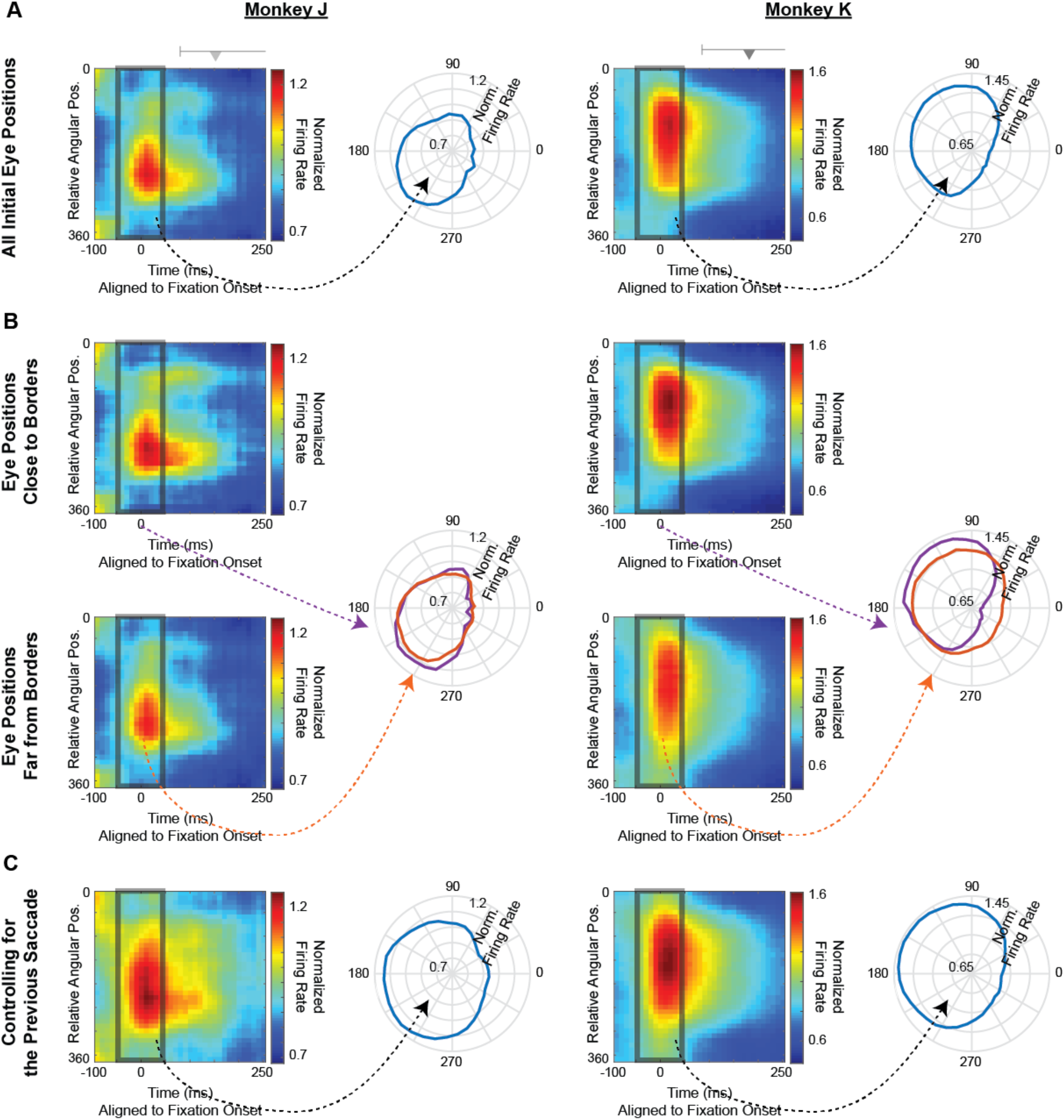
Early population activity reflects the probabilities of upcoming saccades, for individual monkeys. Population activity of Early/Pos neurons as a function of relative angular position, for Monkeys J (left) and K (right). **(A)** On the left, a heat map of normalized activity over time, as a function of relative angular position, averaged across neurons. Ranges of saccade onset times are shown above, as in Fig. 2. On the right, the normalized average activity in the 100 ms surrounding fixation, plotted as a function of the relative angular position. Only saccades away from the PD are included. **(B)** Same as panel A, but now separated for initial eye positions close to the borders (top left, right in purple) and far from the borders (bottom left, right in orange). **(C)** We control for the correlation between the previous and upcoming saccade directions. We only included saccades in which the previous and upcoming saccade directions were less than 90° apart (while in the actual data, previous and upcoming saccades are more likely to be in opposite directions). Unlike in panels A and B, we do not exclude saccades towards the PD.

**Figure S6:**
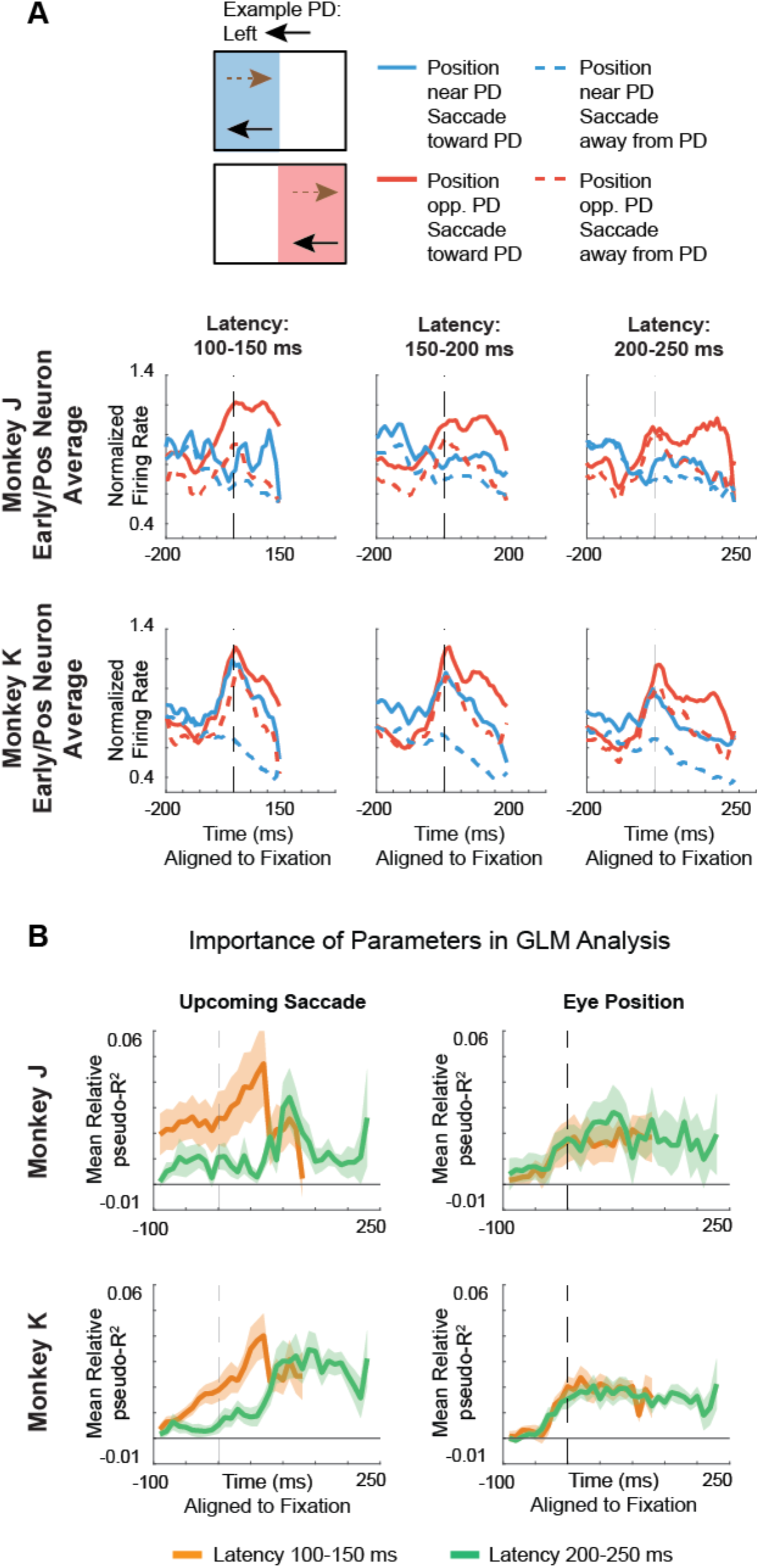
Evolution of activity over time, for individual monkeys. **(A)** PETHs, aligned to fixation onset, of normalized averaged activity of Early/Pos neurons. Blue lines are those with a starting angular eye position near the PD (unlikely that upcoming saccade will be toward PD). Red lines are those with a starting angular eye position opposite the PD (likely that upcoming saccade will be toward PD). Separate PETHs are constructed for saccades with latencies from 100-150 ms (left), 150-200 ms (middle), and 200-250 ms (right). **(B)** Importance of parameters in the generalized linear model, across time, aligned to fixation onset, for Early/Pos neurons. The mean relative pseudo-R^2^ (across Early/Pos neurons) of the upcoming saccade (left) and eye position (right) covariates are shown. We separately determine parameter importance for saccades with latencies of 100-150 ms (orange) and 200-250 ms (green). Shaded areas represent SEMs.

**Figure S7:**
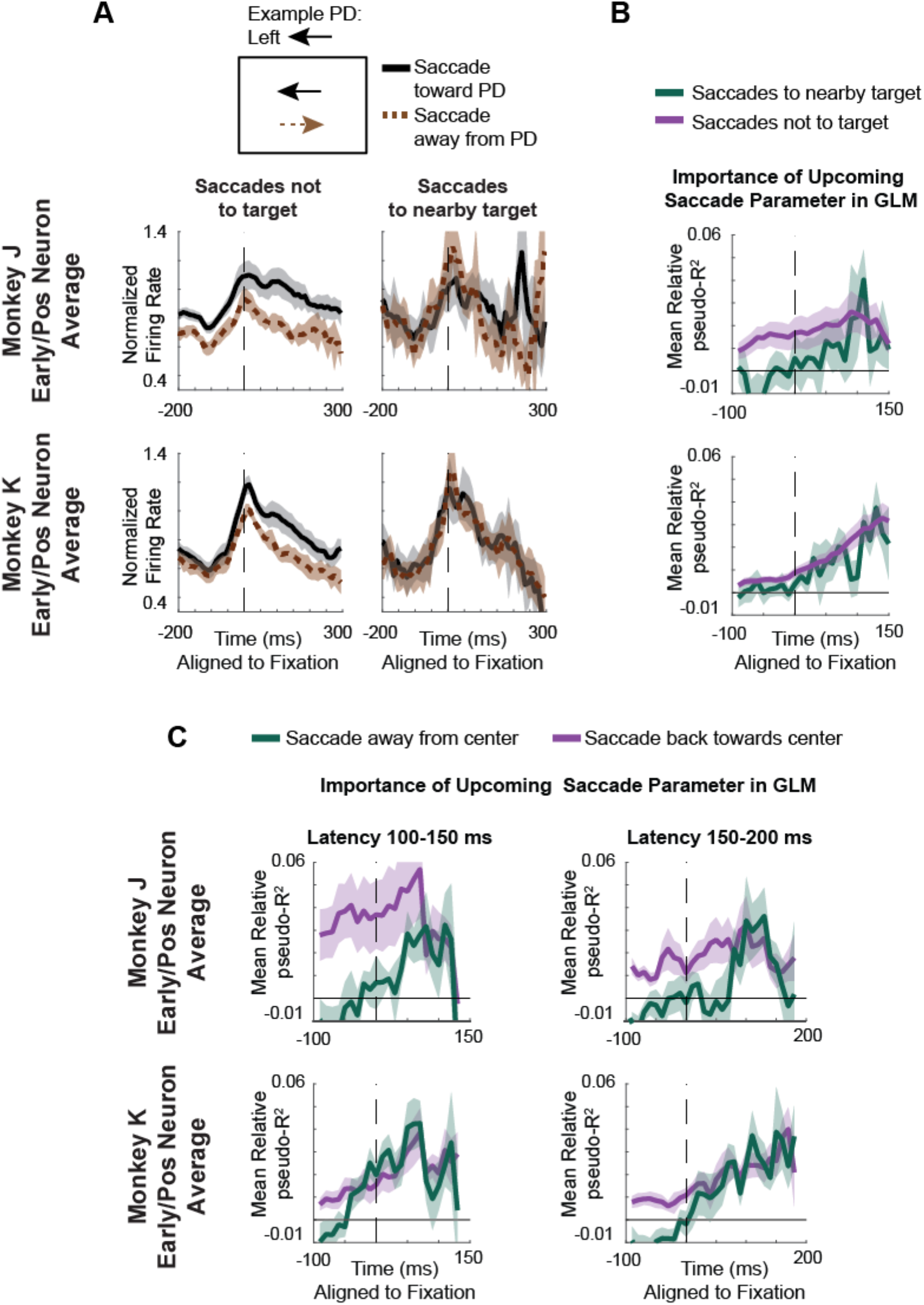
Relationship between early activity and the final selected saccade, for individual monkeys. **(A)** PETHs, aligned to fixation, of saccades toward the preferred direction (PD; black) versus away from the PD (brown, dashed). To control for position, we only use saccades starting from positions opposite the PD. On the left, we only include saccades not to the target. On the right, we only include saccades that go to a nearby target (< 10° away). **(B)** Importance of the upcoming saccade parameter in the generalized linear model, across time, aligned to fixation. The mean relative pseudo-R^2^ (across Early/Pos neurons) was determined for saccades not to the target (purple) and to a target < 10° away (gray). Note that the GLM results were only plot until +150ms, as the results got very noisy since there are limited saccades with latencies > 150 ms. **(C)** Importance of the upcoming saccade parameter in the generalized linear model, across time, aligned to fixation. The mean relative pseudo-R^2^ (across Early/Pos neurons) was determined for saccades back towards the center (purple) and saccades away from the center (gray). Shaded areas represent SEMs. Note that for this comparison, we cannot do a PETH analysis like in previous scenarios, because saccades back to the center, from an angular position opposite the PD, will always be toward the PD (we can’t compare saccades toward versus away from the PD while controlling for position).

**Figure S8:**
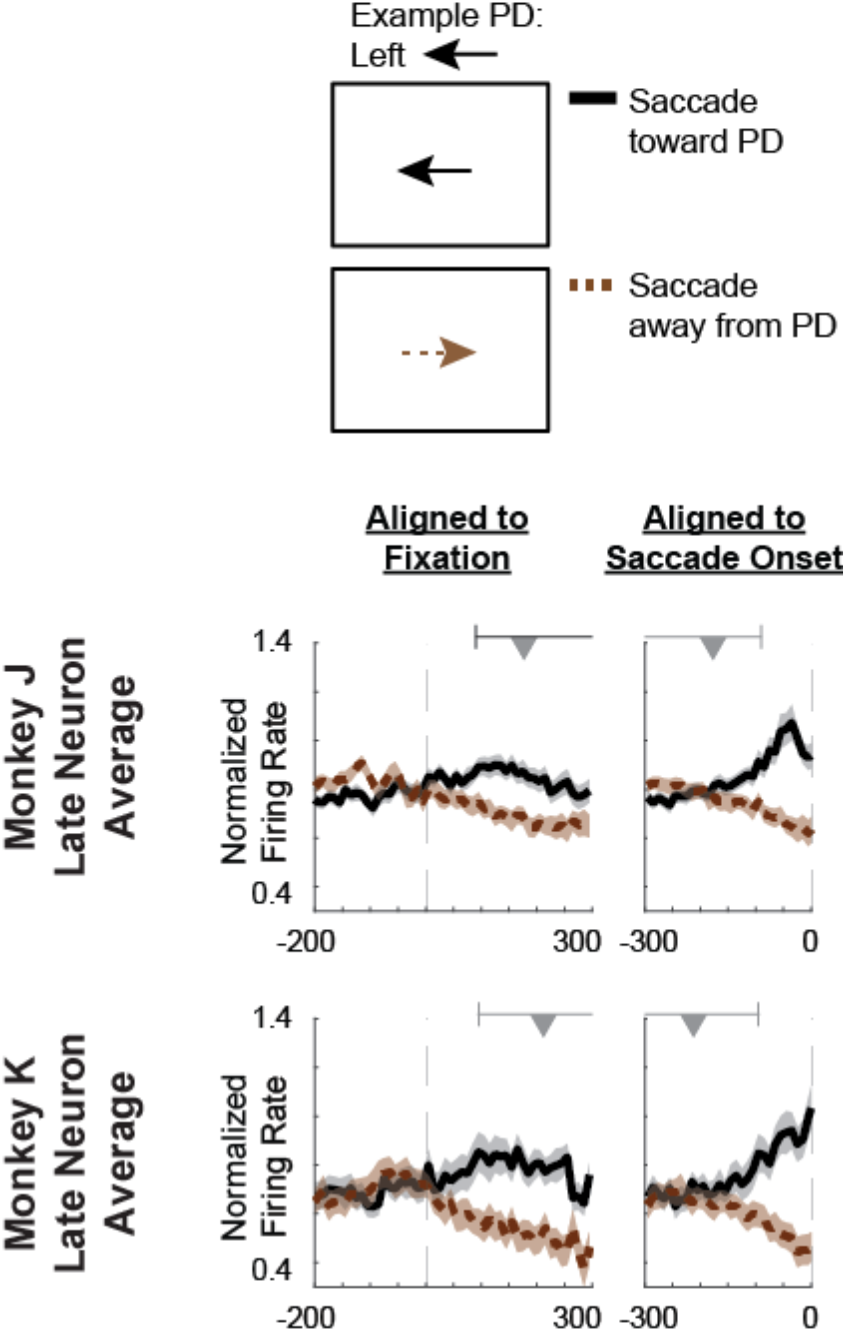
Late Neurons, for individual monkeys. PETHs, aligned both to fixation (left) and the upcoming saccade onset (right), of saccades toward the preferred direction (PD; black) versus away from the PD (brown, dashed). PETHs are averaged across Late neurons. Note that panels B-D of Fig. 7 are not replicated here for individual monkeys, because there was only a single Late/Pos neuron for Monkey K.

**Figure S9:**
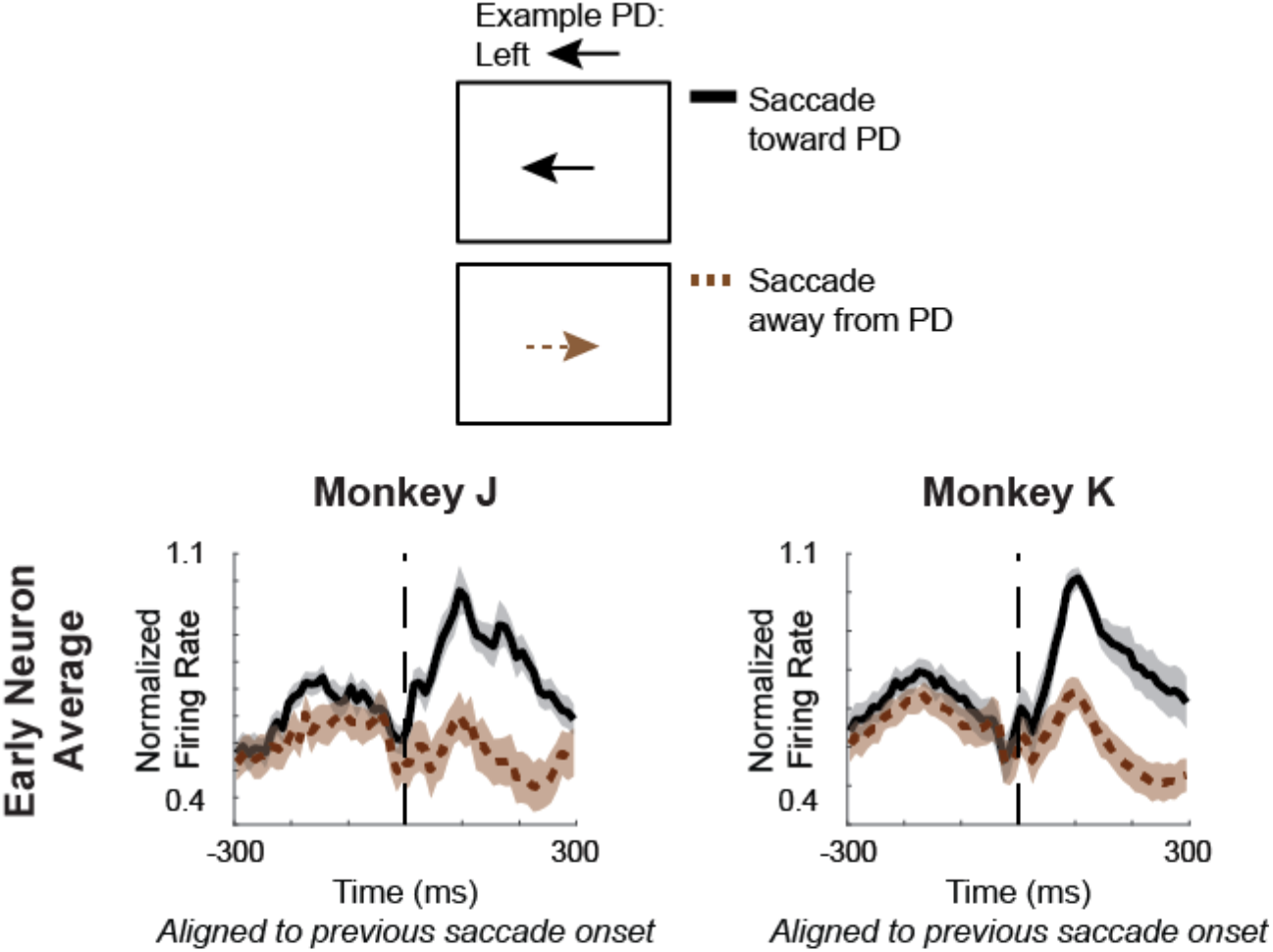
PETHs aligned to the onset of the previous saccade. Peri-event time histograms (PETHs) aligned to the onset of the previous saccade, for saccades toward the preferred direction (PD; black) versus away from the PD (brown, dashed).

**Figure S10:**
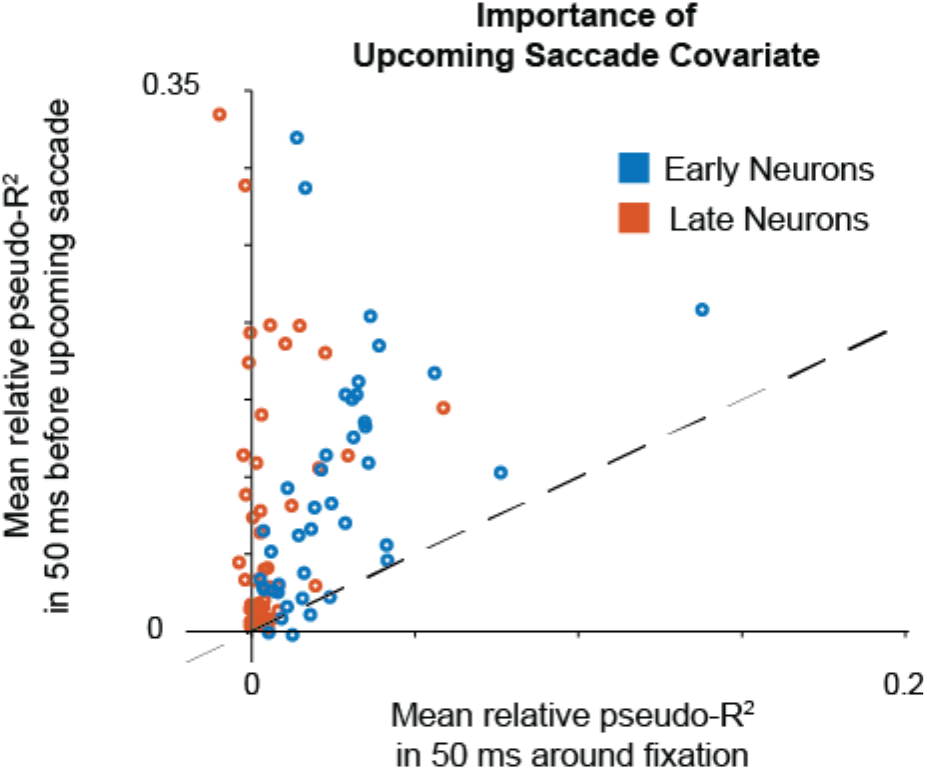
Early and late saccade predictive activity. GLMs were fit with the previous and upcoming saccade as covariates, as done when classifying neurons as “Early” or “Late”. Mean (across cross-validation folds) relative pseudo-R^2^ values of the upcoming saccade covariate are shown when the model is fit in the 50 ms around fixation (x-axis) and the 50 ms before the upcoming saccade (y-axis), for Early neurons (blue) and Late neurons (orange). Dots above/below the dashed line have more/less unique information about the upcoming saccade near the time of the upcoming saccade. Note that the Late neurons with positive x-axis values are not significantly greater than 0.

**Figure S11:**
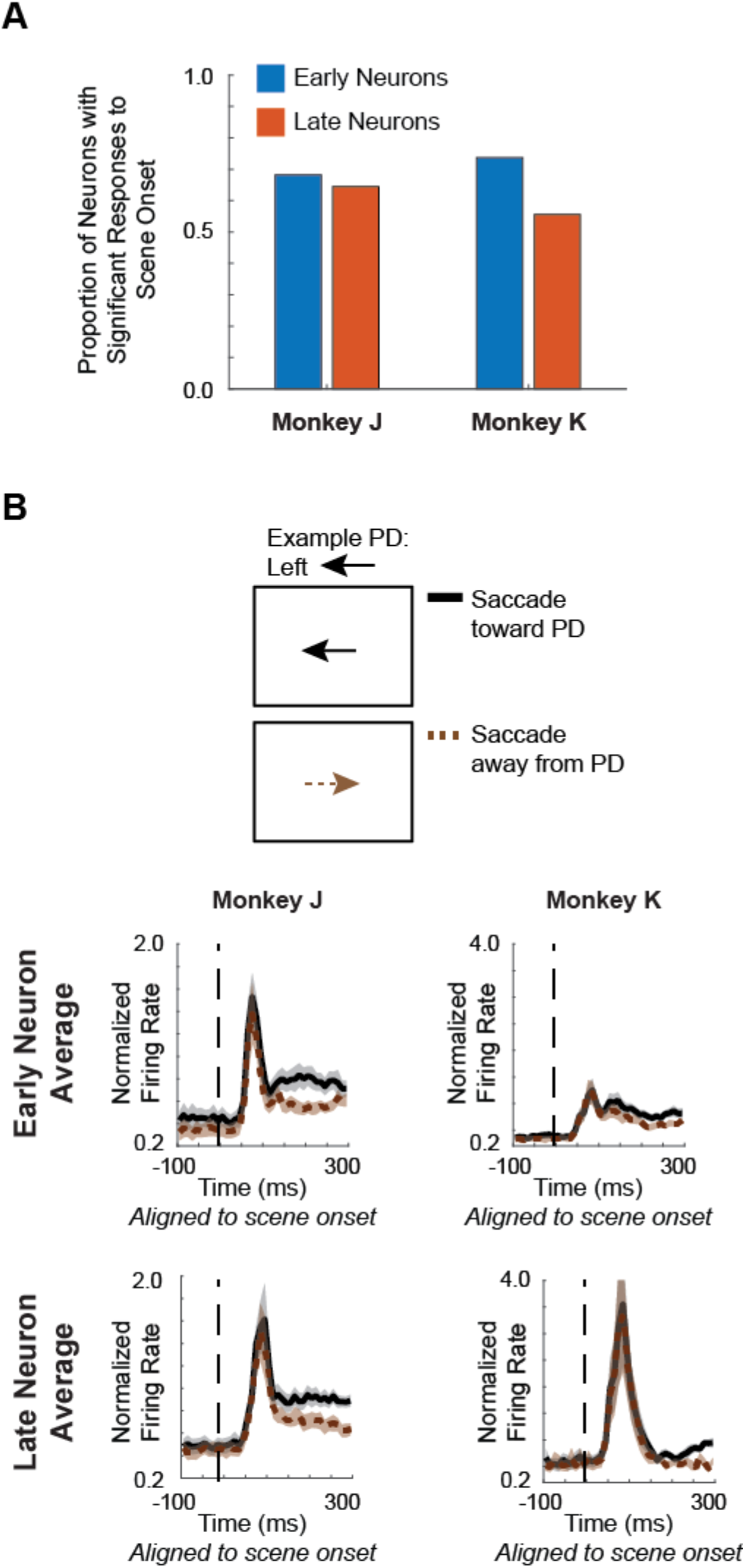
Visual responses of Early and Late neurons. **(A)** The proportion of Early (blue) and Late (orange) neurons that have significant modulation of activity following scene onset (see *Methods*). **(B)** Peri-stimulus time histograms (PSTHs) aligned to the onset of the scene. PSTHs are divided based on whether the upcoming saccade (first saccade of the trial) is toward the preferred direction (PD; black) versus away from the PD (brown, dashed). As opposed to the other PETH figures, here each neuron’s activity is normalized based on the peak average firing rate when making a saccade into its PD. Thus, these plots represent the visual response relative to the movement response.

## References

1. Bichot NP, Schall JD (1999) Effects of similarity and history on neural mechanisms of visual selection. Nature neuroscience 2: 549–554.

2. Bichot NP, Schall JD (2002) Priming in macaque frontal cortex during popout visual search: feature-based facilitation and location-based inhibition of return. The Journal of Neuroscience 22: 4675–4685.

3. Burman DD, Segraves MA (1994) Primate Frontal Eye Field Activity ring Natural Scanning Eye Movements.

4. Phillips AN, Segraves MA (2010) Predictive activity in macaque frontal eye field neurons during natural scene searching. Journal of neurophysiology 103: 1238–1252.

5. Basso MA, Wurtz RH (1998) Modulation of neuronal activity in superior colliculus by changes in target probability. The Journal of Neuroscience 18: 7519–7534.

6. Dorris MC, Munoz DP (1998) Saccadic probability influences motor preparation signals and time to saccadic initiation. The Journal of Neuroscience 18: 7015–7026.

7. Everling S, Munoz DP (2000) Neuronal correlates for preparatory set associated with pro-saccades and anti-saccades in the primate frontal eye field. Journal of Neuroscience 20: 387–400.

8. Mirpour K, Arcizet F, Ong WS, Bisley JW (2009) Been there, seen that: a neural mechanism for performing efficient visual search. Journal of Neurophysiology 102: 3481–3491.

9. Everling S, Dorris MC, Klein RM, Munoz DP (1999) Role of primate superior colliculus in preparation and execution of anti-saccades and pro-saccades. Journal of Neuroscience 19: 2740–2754.

10. Zhou H, Desimone R (2011) Feature-based attention in the frontal eye field and area V4 during visual search. Neuron 70: 1205–1217.

11. Paré M, Munoz DP (2001) Expression of a re-centering bias in saccade regulation by superior colliculus neurons. Experimental Brain Research 137: 354–368.

12. Glaser JI, Wood DK, Lawlor PN, Ramkumar P, Kording KP, et al. (2016) The role of expected reward in frontal eye field during natural scene search. Journal of neurophysiology: jn. 00119.02016.

13. Ramkumar P, Lawlor PN, Glaser JI, Wood DK, Phillips AN, et al. (2016) Feature-based attention and spatial selection in frontal eye fields during natural scene search. Journal of neurophysiology: jn. 01044.02015.

14. Tseng P-H, Carmi R, Cameron IG, Munoz DP, Itti L (2009) Quantifying center bias of observers in free viewing of dynamic natural scenes. Journal of vision 9: 4–4.

15. Bindemann M (2010) Scene and screen center bias early eye movements in scene viewing. Vision research 50: 2577–2587.

16. Buswell GT (1935) How people look at pictures: a study of the psychology and perception in art.

17. Zambarbieri D, Beltrami G, Versino M (1995) Saccade latency toward auditory targets depends on the relative position of the sound source with respect to the eyes. Vision research 35: 3305–3312.

18. Fuller JH (1996) Eye position and target amplitude effects on human visual saccadic latencies. Experimental Brain Research 109: 457–466.

19. McPeek RM, Keller EL (2004) Deficits in saccade target selection after inactivation of superior colliculus. Nature neuroscience 7: 757–763.

20. Hikosaka O, Wurtz R (1986) Saccadic eye movements following injection of lidocaine into the superior colliculus. Experimental Brain Research 61: 531–539.

21. Dorris MC, Pare M, Munoz DP (1997) Neuronal activity in monkey superior colliculus related to the initiation of saccadic eye movements. Journal of Neuroscience 17: 8566–8579.

22. Rivaud S, Müri R, Gaymard B, Vermersch A, Pierrot-Deseilligny C (1994) Eye movement disorders after frontal eye field lesions in humans. Experimental Brain Research 102: 110–120.

23. Colby C, Goldberg M (1992) The updating of the representation of visual space in parietal cortex by intended eye movements. Science 255: 90–92.

24. Sommer MA, Wurtz RH (2006) Influence of the thalamus on spatial visual processing in frontal cortex. Nature 444: 374.

25. Zirnsak M, Steinmetz NA, Noudoost B, Xu KZ, Moore T (2014) Visual space is compressed in prefrontal cortex before eye movements. Nature 507: 504.

26. Fernandes HL, Stevenson IH, Phillips AN, Segraves MA, Kording KP (2013) Saliency and saccade encoding in the frontal eye field during natural scene search. Cerebral Cortex 24: 3232–3245.

27. Churan J, Guitton D, Pack CC (2011) Context dependence of receptive field remapping in superior colliculus. Journal of neurophysiology 106: 1862–1874.

28. Walker MF, Fitzgibbon EJ, Goldberg ME (1995) Neurons in the monkey superior colliculus predict the visual result of impending saccadic eye movements. Journal of neurophysiology 73: 1988–2003.

29. Russo GS, Bruce CJ (1993) Effect of eye position within the orbit on electrically elicited saccadic eye movements: a comparison of the macaque monkey’s frontal and supplementary eye fields. Journal of neurophysiology 69: 800–818.

30. Cassanello CR, Ferrera VP (2007) Computing vector differences using a gain field - like mechanism in monkey frontal eye field. The Journal of physiology 582: 647–664.

31. Gold JI, Shadlen MN (2000) Representation of a perceptual decision in developing oculomotor commands. Nature 404: 390–394.

32. Ding L, Gold JI (2011) Neural correlates of perceptual decision making before, during, and after decision commitment in monkey frontal eye field. Cerebral Cortex: bhr178.

33. Campos M, Cherian A, Segraves MA (2006) Effects of eye position upon activity of neurons in macaque superior colliculus. Journal of neurophysiology 95: 505–526.

34. Van Opstal A, Hepp K, Suzuki Y, Henn V (1995) Influence of eye position on activity in monkey superior colliculus. Journal of Neurophysiology 74: 1593–1610.

35. Sommer MA, Wurtz RH (2002) A pathway in primate brain for internal monitoring of movements. Science 295: 1480–1482.

36. Sommer MA, Wurtz RH (2004) What the brain stem tells the frontal cortex. II. Role of the SC-MD-FEF pathway in corollary discharge. Journal of neurophysiology 91: 1403–1423.

37. Wang X, Zhang M, Cohen IS, Goldberg ME (2007) The proprioceptive representation of eye position in monkey primary somatosensory cortex. Nature neuroscience 10: 640.

38. Stanton GB, Friedman HR, Dias EC, Bruce CJ (2005) Cortical afferents to the smooth-pursuit region of the macaque monkey’s frontal eye field. Experimental brain research 165: 179–192.

39. Sommer MA, Wurtz RH (2000) Composition and topographic organization of signals sent from the frontal eye field to the superior colliculus. Journal of Neurophysiology 83: 1979–2001.

40. Bruce CJ, Goldberg ME (1985) Primate frontal eye fields. I. Single neurons discharging before saccades. J Neurophysiol 53: 603–635.

41. Lowe K, Schall JD (2017) Functional categories in macaque frontal eye field. bioRxiv: 212589.

42. Sajad A, Sadeh M, Yan X, Wang H, Crawford JD (2016) Transition from target to gaze coding in primate frontal eye field during memory delay and memory–motor transformation. eNeuro 3: ENEURO. 00400016.2016.

43. Basso MA, Wurtz RH (1997) Modulation of neuronal activity by target uncertainty. Nature 389: 66–69.

44. Yang T, Shadlen MN (2007) Probabilistic reasoning by neurons. Nature 447: 1075–1080.

45. Glaser JI, Perich MG, Ramkumar P, Miller LE, Kording KP (2017) Population Coding Of Conditional Probability Distributions In Dorsal Premotor Cortex. bioRxiv: 137026.

46. Körding KP, Wolpert DM (2004) Bayesian integration in sensorimotor learning. Nature 427: 244.

47. Jazayeri M, Shadlen MN (2010) Temporal context calibrates interval timing. Nature neuroscience 13: 1020.

48. Darlington TR, Beck JM, Lisberger SG (2018) Neural implementation of Bayesian inference in a sensorimotor behavior. Nature neuroscience 21: 1442.

49. Kim B, Basso MA (2010) A probabilistic strategy for understanding action selection. The Journal of Neuroscience 30: 2340–2355.

50. Bahill AT, Clark MR, Stark L (1975) The main sequence, a tool for studying human eye movements. Mathematical Biosciences 24: 191–204.

51. Waldhör T, Haidinger G, Schober E (1998) Comparison of R2 measures for Poisson regression by simulation. Journal of Epidemiology and Biostatistics 3: 209–215.

52. Fernandes HL, Stevenson IH, Phillips AN, Segraves MA, Kording KP (2013) Saliency and saccade encoding in the frontal eye field during natural scene search. Cerebral Cortex: bht179.

